# IDENTIFICATION AND QUANTIFICATION OF MATRISOME PROTEINS OF MOUSE KIDNEYS

**DOI:** 10.1101/2022.07.06.499068

**Authors:** Umut Rende, Seong Beom Ahn, Subash Adhikari, Anna Guller

**Affiliations:** ARC Centre of Excellence in Nanoscale Biophotonics, The Graduate School of Biomedical Engineering, University of New South Wales, Sydney, NSW 2052, Australia; Macquarie Medical School, Macquarie University, NSW 2109, Australia; The Walter and Eliza Hall Institute of Medical Research, Parkville, Victoria 3052, Australia; Biomolecular Discovery Research Centre, Macquarie University, NSW 2109, Australia

**Keywords:** Extracellular matrix, matrisome, kidneys, mouse, tissue extraction, proteomics, mass spectrometry

## Abstract

Extracellular matrix (ECM) is essential for tissue homeostasis. Understanding the matrisome (ECM proteome) composition and mechanisms of ECM control in health and disease is crucial for discovering therapeutic agents and diagnostic tools for inflammatory, fibrotic and cancerous conditions. The challenging obstacle in the ECM analysis is the need to optimise matrisome enrichment methods for different organs, diseases, and species. Currently, there is no optimized protocol nor a publicly available matrisome database for mouse kidneys. This limits the power of murine models in renal diseases and development research. In this study, we comparatively explored the matrisome of healthy C57BL/6 mice using two matrisome extraction methods, including the Millipore Compartment Fractionation (Method-1) and the Sequential Extraction (Method-2) approaches. We examined the efficiency of these methods in matrisome profiling by LC-MS/MS, protein identification and label-free quantification using MaxQuant. As a result of the study, 113 matrisome proteins were identified, including 22 proteins that have not been previously listed in the Matrisome Database (MD). Method-2 allowed identification and quantification of all core and ECM-associated matrisome proteins detected by Method-1 and additionally revealed more core matrisome and ECM-associated proteins. By characterisation of the murine renal matrisome enhanced by our methodological insights, this study provides critically important information for biological and medical kidney research.

## INTRODUCTION

Extracellular matrix (ECM) is a complex meshwork of multiple components such as structural proteins, glycoproteins (GPs), polysaccharides, proteoglycans (PGs), and ECM-remodelling enzymes, as well as ECM receptors, growth factors and cytokines. ECM provides adhesion and anchorage to the cells ^1^, plays a role in tissue & organ development ^2^, cell signalling and cell metabolism ^3^, wound healing and disease responses ^4^. Impairment in the function of ECM proteins, including excessive deposition or destruction, habeen linked to many diseases ^5^. Thus, better understanding nature of ECM proteins and its homeostasis can illuminate many underlying pathophysiological events.

An important aspect of the ECM proteome, which is also known as matrisome, is defining matrisome proteins. *In silico*, matrisome is defined in two divisions by Naba *et al* ^6^; structural ECM components (core matrisome) and ECM interacting components (ECM-associated matrisome) based on the presence of signal peptide, ECM protein domains, ECM interacting domains and/or remodelling ECM. The core matrisome constitutes categories of collagens, ECM glycoproteins and proteoglycans and characterized as extensively glycosylated, having multidomains and expanded shapes which favors in forming fiber and/or supramoleculars ^7,8^, while excluding proteins which have also transmembrane, tyrosine kinase and phosphatase domains such as growth factor receptors and integrins ^9^. On the other hand, proteins which were found in ECM, but could not be categorized as core matrisome are defined as ECM-associated matrisome molecules ^9^. This division comprises categories of a) ECM-affiliated proteins, which has similar architecture with ECM proteins and/or known to be associated with ECM proteins, b) ECM regulators (ECM-remodeling enzymes, crosslinkers, proteases, regulators) and c) secreted factors which are known interact with ECM ^9,10^. This classification remains open for debate and additional ECM protein discoveries will lead to a better understanding of the ECM proteome. The matrisome proteins identified in different tissues and tumors of both human and mouse origin are deposited in “Matrisome Database (MD)” (http://matrisomeproject.mit.edu/).

The analysis of composition of ECM in normal and pathological conditions using mass spectrometry-based proteomics is urged in the last decade. Proteomics provide information regarding protein synthesis, degradation, modifications and also protein interactions with other molecules. Thus, mass spectrometry based-proteomics has being used largely for discovery of biomarkers and diagnostic tools ^11^. Rapid technological developments in proteomics such as enhanced sample preparation protocols, improved capabilities in mass spectrometry (MS), database searching, and bioinformatics analysis allow unbiased protein quantification, identification and characterization including discovery of post-translational modifications (PTMs) ^12^.

In spite of developments in proteomics field, standard proteomic techniques, which uses whole tissue lysate, cannot provide a comprehensive information about crosslinked and highly insoluble matrix elements. Some of the challenges in ECM proteomics were to separate and identify low abundant matrisome proteins from intracellular proteins, to solubilize heavily cross-linked ECM proteins and the presence of abundant PTMs which affected the separation and identification ^13^.

Recent advancements in matrisome protein extractions and separations provided subsequent analysis with mass spectrometry. For example, one of the recently developed extraction methods is the compartmental protein fractionation approach ^6,14–17^. This approach involves sequential incubations in buffers of different pH and salt & detergent concentrations which results in biochemical separation of cytosolic, nuclear, membrane and cytoskeletal proteins and the enrichment of matrisome proteins. Another developed method is using guanidine hydrochloride (GuHCl) for the solubilization of insoluble matrisome proteins after a decellularization step ^17–20^. This extraction method is performed using an ionic (NaCl) buffer to extract loosely bound ECM proteins (enzymes, secreted factors, ECM-associated and newly deposited proteins), as well as low detergent (sodium dodecyl sulphate, SDS) concentration with shorter incubation to eliminate intracellular proteins and GuHCl buffer to obtain heavily crosslinked ECM proteins.

In biomedical research, mice have a critical role to study organ development, mechanisms of diseases and to discover new drug targets and biomarkers since murine models are reasonably easy to maintain, reproduce rapidly and cost less^21,22^. For example, mouse models are essential for understanding of tubulointerstitial fibrosis, described as excessive accumulation of ECM components and a key pathological feature of chronic kidney diseases (CKDs) leading to renal failure ^23^. However, no optimized protocol, nor a publicly available matrisome database are currently available for mouse kidneys. This limits the progress in ECM related kidney disease research. Recently, McCabe *et al* 2021 ^17^ and Lipp *et al* 2021 ^24^ extracted mouse kidney matrisome proteins using a compartment protein fractionation approach and identified ECM proteins in the insoluble fractions. However, in these studies, identification of loosely bound ECM proteins and quantification of identified ECM proteins have not been not performed.

Here, we aimed to identify matrisome proteins and compare their availability and abundances using two widely applied extraction methods in healthy kidneys of C57BL/6 mice. We used compartmental protein fractionation (further labelles as Method 1) and sequential extraction using GuHCl (labelled as Method 2) approaches, and the extracted matrisome proteins were analysed with LC-MS/MS system. To classify matrisome proteins, the identified proteins were analysed with the Matrisome Database (MD) and UniProt databases. To compare the abundance of identified matrisome proteins from the two extraction methods, MaxQuant label-free quantification (LFQ) was employed and then the data was analyzed using LFQ-Analyst^25^. The project workflow is depicted in **Figure 1**.

**Figure 1.**
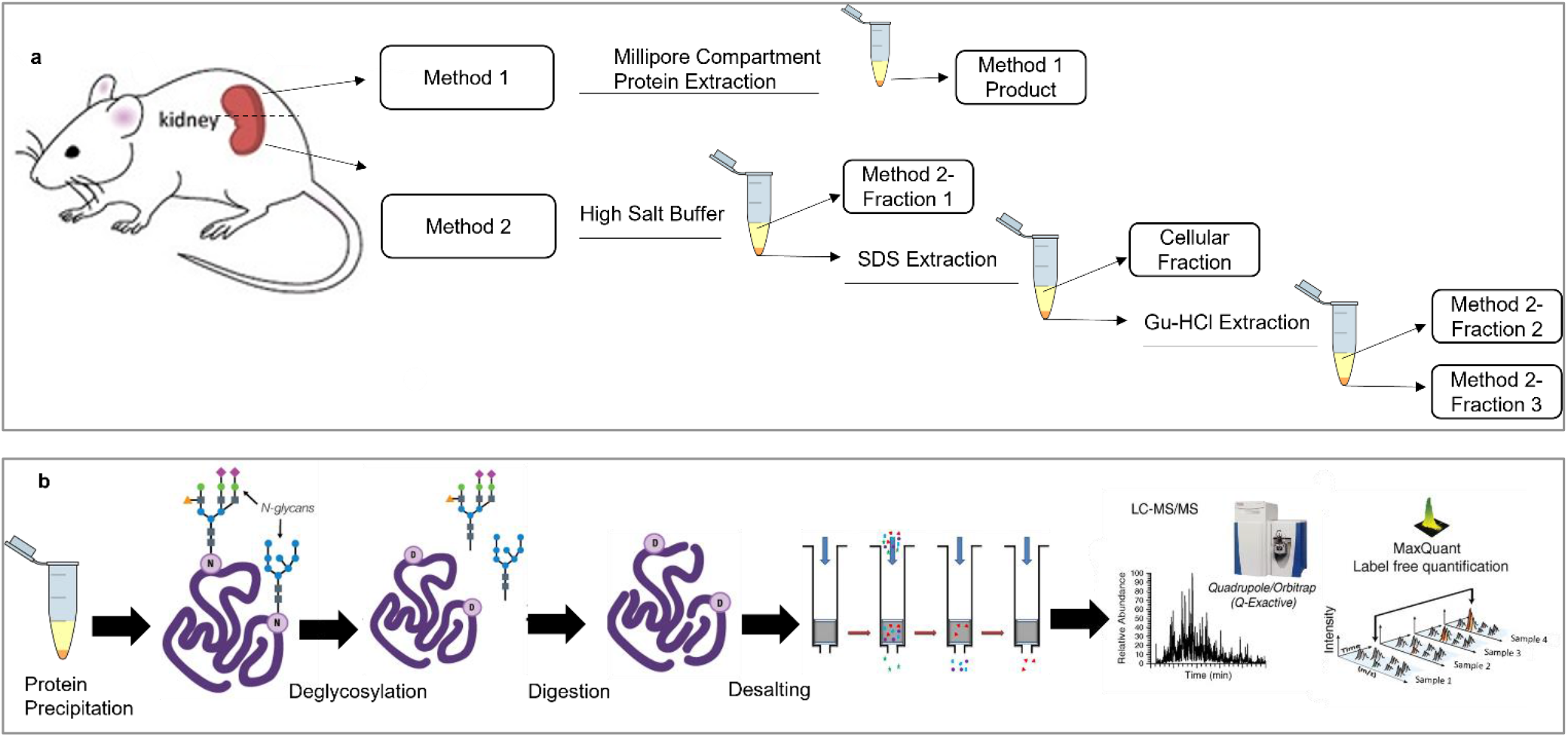
Schematic illustration of the methods applied in the current study. (a) Illustrates matrisome protein extraction by Method 1 and 2 from healthy mouse kidneys. (b) Illustrates the further processes of proteins after obtaining samples from Method 1 and 2. SDS (Sodium Dodecyl Sulfate) and Gu-HCl (Guanidine Hydrochloride).

## RESULTS

### Qualitative analysis of mouse kidney matrisome

#### Identification of the proteins of the mouse kidney matrisome

In total, 2442 unique peptides were identified using Methods 1 and 2. Collectively, this resulted in a total of 113 matrisome protein identifications in the healthy mouse kidneys (**Table S1**).

The overview of the mouse matrisome composition identified in the current study is shown in **Table 1**. Among the identified proteins, 51 (45%) were classified as core matrisome proteins. The rest 62 identified proteins (55%) were accounted as matrisome-associated proteins. The core matrisome of healthy mouse kidneys was dominated by ECM glycoproteins, followed by the collagens and ECM proteoglycans (**Table 1**, Ratio, % of the division). The most frequently identified matrisome-associated proteins belonged to the category of ECM regulators, followed by the ECM-affiliated proteins and secreted factors.

**Table 1.** Composition of mouse kidney matrisome proteins identified in this study.

| Matrisome Division | Matrisome Category | N of identified proteins | Ratio, % of the division | Ratio, % of the whole matrisome |
| --- | --- | --- | --- | --- |
| Core matrisome | ECM Glycoproteins | 28 | 55 | 25 |
|  | Collagens | 16 | 31 | 14 |
|  | Proteoglycans | 7 | 14 | 6 |
| Matrisome-associated proteins | ECM Regulators | 36 | 58 | 32 |
|  | ECM-affiliated Proteins | 19 | 31 | 17 |
|  | Secreted Factors | 7 | 11 | 6 |

Notably, 22 proteins among the 113 identified ECM proteins have not been previously classified in the MD (**Table 2**). These 22 proteins were attributed as the matrisome components based on their extracellular location, functions, and also on their interactions with extracellular proteins and were classified into the Matrisome category (the relevant references are provided in Table 2). The majority of the newly classified mouse kidney matrisome proteins belonges to the division of the matrisome-associated proteins, including 14 ECM regulators, 5 ECM-affiliated proteins, and 2 secreted factors, and only one was classified as a core matrisome component (an ECM glycoprotein). Interestingly, 6 of these matrisome proteins identified in mouse kidneys for the first time were revealed only by Method 2, emphasizing its advantages in the ECM proteins’ enrichment.

**Table 2.**
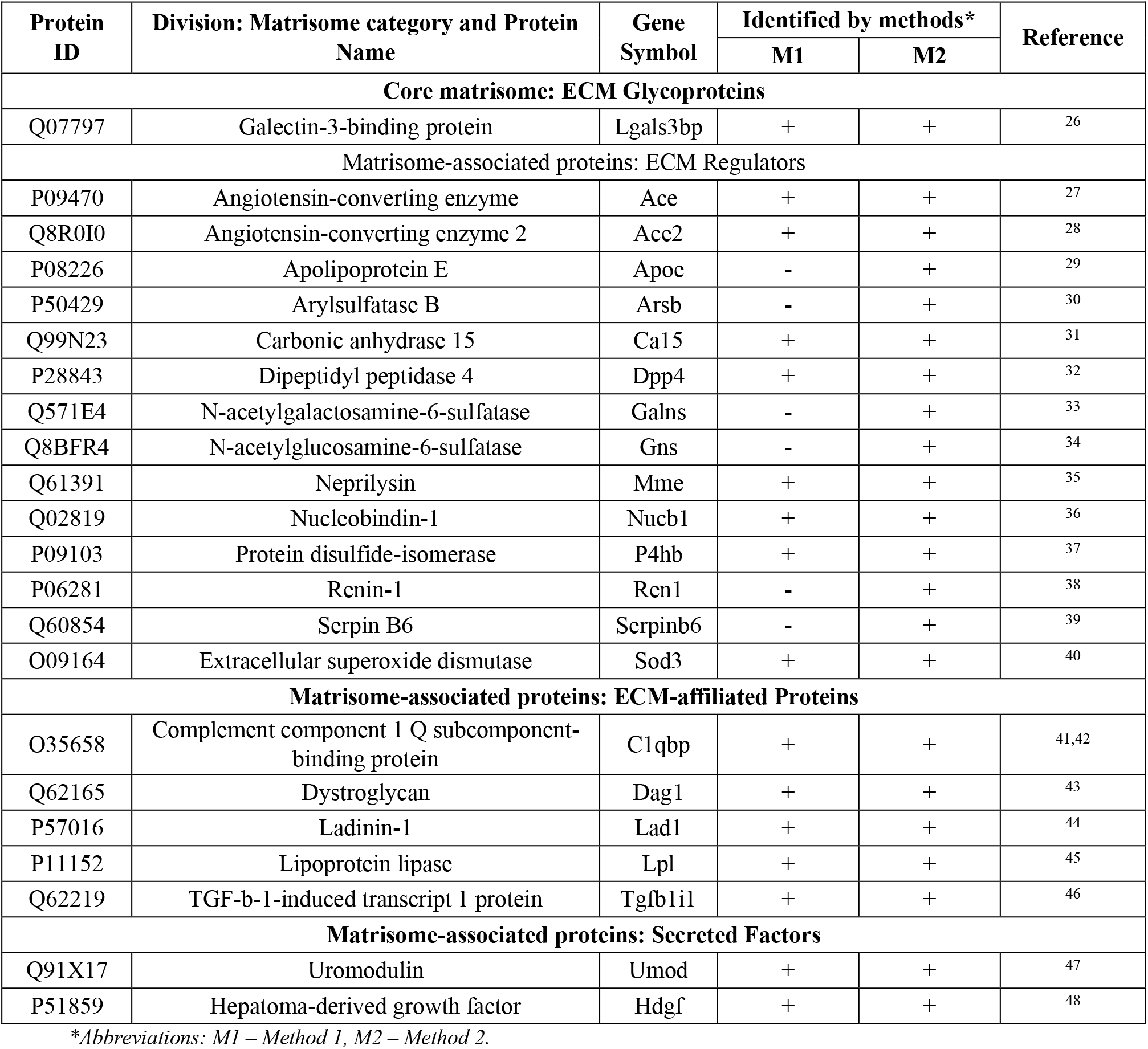
Newly detected and classified mouse kidneys matrisome proteins that are not listed in the curent Matrisome Database.

#### Gene ontology analysis of mouse matrisome proteins

Gene Ontology (GO) analysis based on biological processes annotated 100 mouse matrisome proteins (from 113 proteins) (**Figure 2**). The annotation showed that the majority of the identified matrisome proteins shared the relationship to the class of “cellular process” (89/100), followed by “biological regulation” (83/100) and “developmental process” (63/100) (**Figure 2a**). The rest of the proteins belonged to classes of response to stimulus (54/100), metabolic process (45/100), multicellular organismal process (35/100), biological adhesion (34/100), localization (27/100), immune system process (17/100), locomotion (17/100), biological process involved in interspecies (17/100), reproductive process (14/100), behavior (10/100), biomineral tissue development (3/100), growth (3/100), viral process (2/100), trans-synaptic signaling (2/100), removal of superoxide radicals (1/100) and estrous cycle (1/100). These attributions indicate massive involvement of the matrisome in the regulation of the cellular physiological activity in the kidneys.

**Figure 2.**
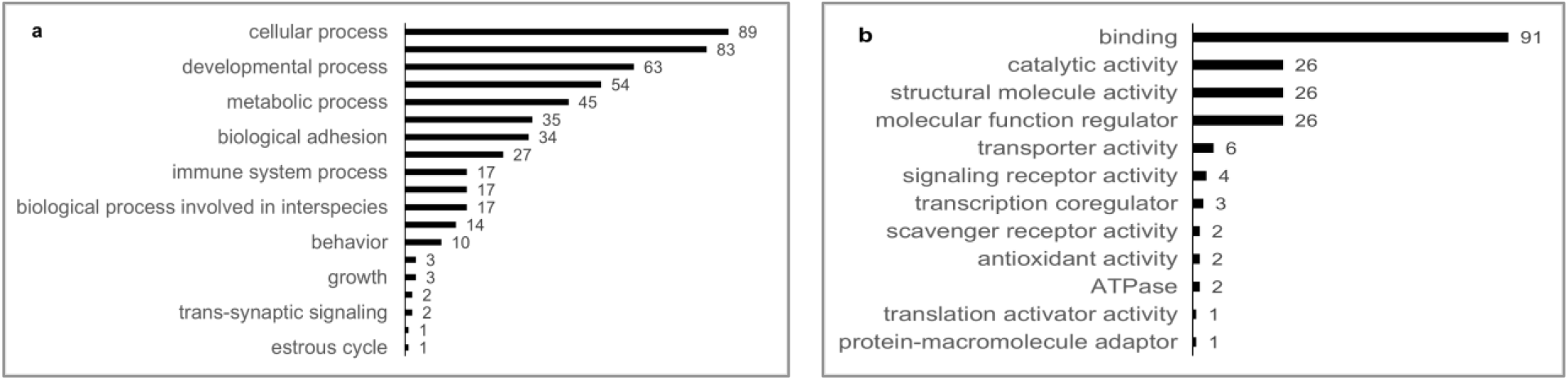
Classification of majority of the identified mouse kidney matrisome proteins by GO terms according to the UniProt Database: **(a)** GO terms for the Biological Process and **(b)** GO terms for the Molecular Function.

GO analysis by terms based on the molecular function of proteins (annotated 106 proteins from 113) revealed that the majority of (93/106) of identified proteins as “binding” proteins which interact with other molecules with a specific site. This was followed by “catalytic activity” (26/106), “structural molecule activity” (26/106) and “molecular function regulator (26/106)” (**Figure 2b**). The rest of the proteins belonged to classes of transporter activity (6/106), signaling receptor activity (4/106), transcription coregulator (3/106), scavenger receptor activity (2/106), antioxidant activity (2/106), ATPase (2/106), translation activator activity (1/106) and protein-macromolecule adaptor (1/106).

#### The comparative efficiency of different extraction methods in detection of matrisome proteins

To define which method is more efficient in revealing the matrisome proteins in mouse kidney tissues, we compared list of the matrisome proteins obtained from Methods 1 and 2 and classified them into matrisome categories. This comparison is visualized in **Figure 3**.

**Figure 3.**
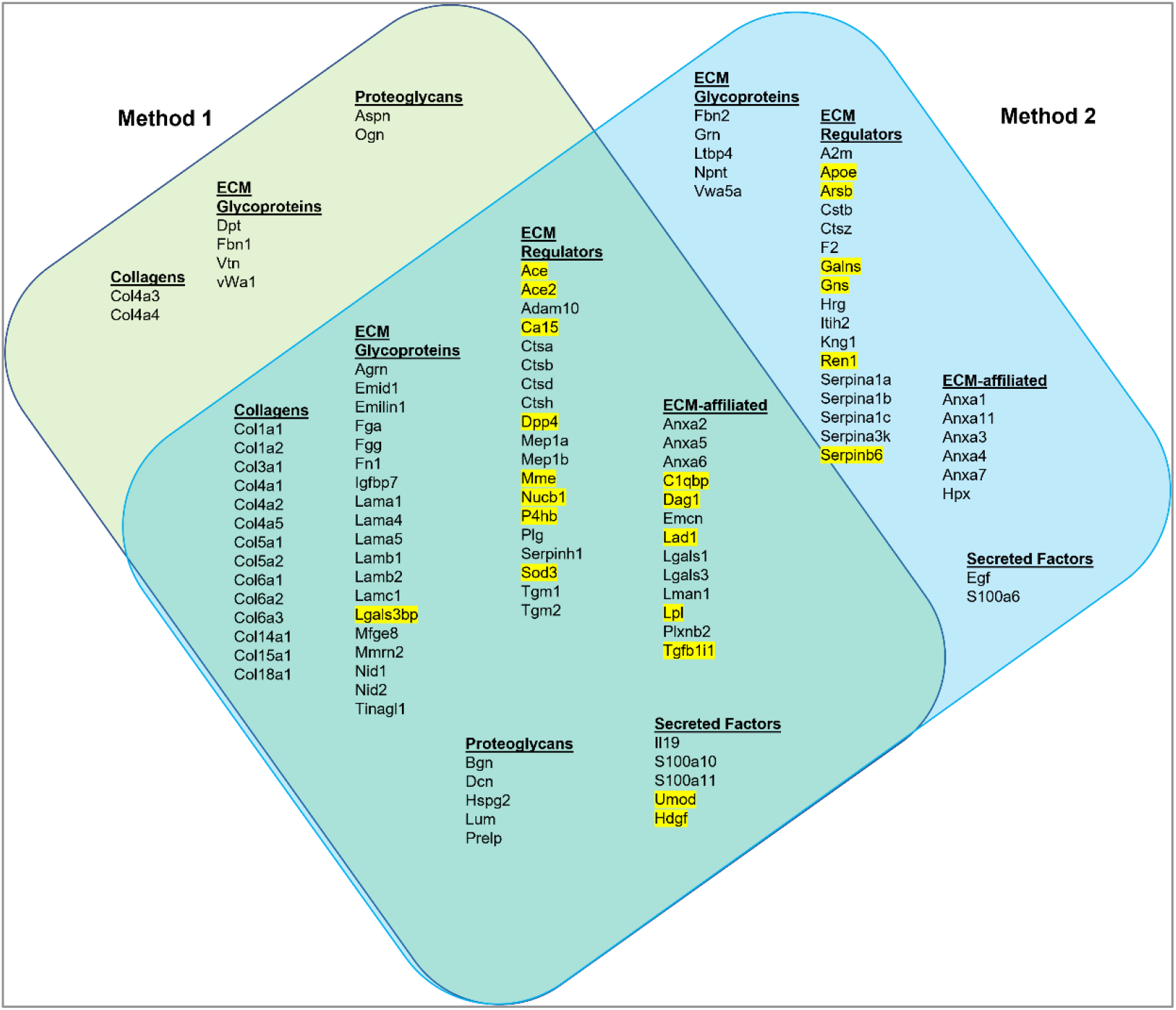
Identified matrisome proteins with unique peptides from Methods 1 and 2. Venn diagram illustrates the common and unique matrisome proteins in each method. Matrisome proteins were classified as Collagens, ECM Glycoproteins, Proteoglycans, ECM-affiliated, ECM Regulators and Secreted Factors. The proteins highlighted proteins in yellow were not previously listed in the MD.

As shown in **Figure 3**, from the 113 matrisome proteins, Method 1 was able to identify 83 proteins whereas Method 2 revealed 105 proteins (73.5% vs 92.9%; CI_95_% [64.4; 82.5]% vs [87.3; 98.5]%, respectively). This indicates Method 2 has a statistically significantly higher efficiency of matrisome protein enrichment/identification compared to Method 1. It is also visible that Method 2 was particularly more efficient than Method 1 in the identification of ECM regulators.

Seventy five matrisome proteins were detected by both methods, 8 proteins were uniquely detected in Method 1 (2 collagens, 4 ECM glycoproteins and 2 proteoglycans) and 30 proteins were only detected in Method 2 (5 ECM glycoproteins, 17 ECM regulators, 6 ECM-affiliated proteins and 2 secreted factors). Additonal comparison of the results obtained in mouse kidney matrisome proteins identification vs the Matrisome Database divisions and categories can be found in **Figure S1** and **Table S1** in Supporting Information.

#### Comparative analysis of the composition of the mouse and human kidney matrisome

We also compared the mouse matrisome proteins (identified in healthy mouse kidneys in our and other mouse kidney studies ^17,24,49^) with the published data on the matrisome proteins from healthy human kidneys ^50,51^. We extracted data across currently available studies where different protein extraction and analysis methods were used. A combined list of matrisome proteins identified in mouse and human kidneys is shown in **Table S2.** The comparative analysis of the number of proteins comprising individual matrisome categories of human and mouse kidneys according to our and literature data is presented in **Table S3** in Supporting Information.

By combining our data with other published data obtained in mouse kidney matrisome studies ^17,49,52^, 229 mouse kidney matrisome proteins were identified (see **Table S2**). On the other hand, recent studies ^53,54^ together identified 178 matrisome proteins in adult human kidneys. The comparison of mouse and human data revealed that 134 matrisome proteins that were shared between mouse and human kidney matrisomes, while 95 of these proteins were found only in mouse kidneys and 44 were identified only in human-specific (**Table S2**). Across these studies, 20 proteins (10 ECM Regulators, 6 ECM affiliated and 4 Secreted Factors) could be identified as matrisome components *de novo* in our study.

### Mouse Kidney Matrisome Protein Quantification

The quantitative characterization of healthy mouse kidney matrisome was performed by analysing the MaxQuant LFQ protein intensities data. The analysis revealed a total of 87 distinct matrisome proteins that were able to be quantitatively examined (**Table S4** in Supplementary Information).

From the total LFQ intensities, the core matrisome of the healthy mouse kidneys is predominantly composed by ECM glycoproteins (45% or 51%, as quantified by Method 1 and 2, respectively). Collagens comprise 36% of the core mouse kidney matrisome for both Methods. Proteoglycans form 19% or 13% of the core matrisome, according to the quantifications of the samples prepared by Method 1 and 2, respectively (**Figures 4a** and **4b**). The matrisome-associated proteins in mouse kidneys are presented mostly by the ECM regulators, followed by the ECM-affiliated proteins and secreted factors (**Figures 4c** and **4d**). The detailed composition of the quantified proteins in mouse kidney matrisome is visualized in **Figure 5.**

**Figure 4.**
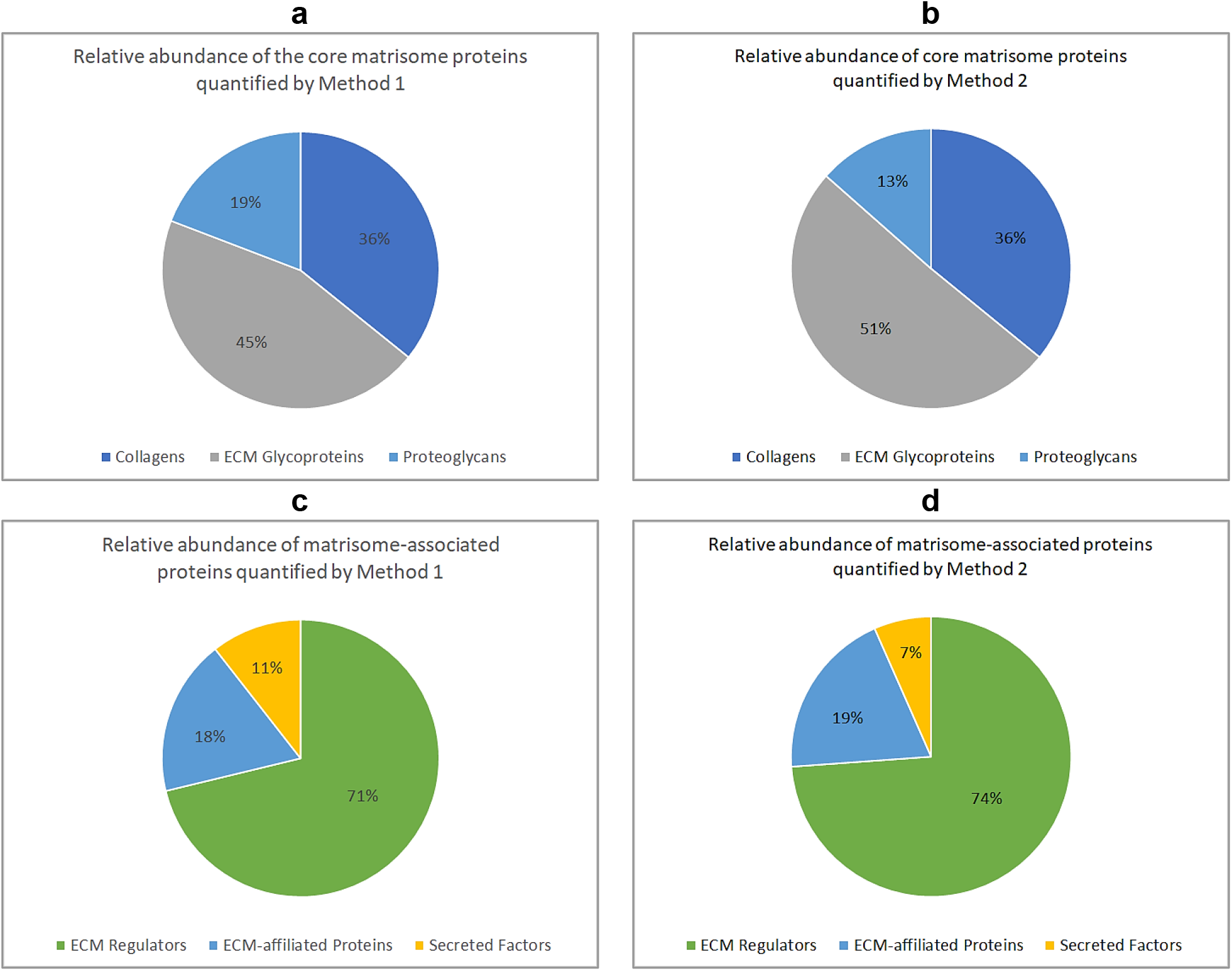
Relative abundance of matrisome proteins in healthy mouse kidneys according to MaxQuant LFQ protein intensities data. Core matrisome proteins quantified by Method 1 **(a)** and 2 **(b).** Matrisome-associated proteins quantified by Method 1 **(c)** and 2 **(d).**

**Figure 5.**
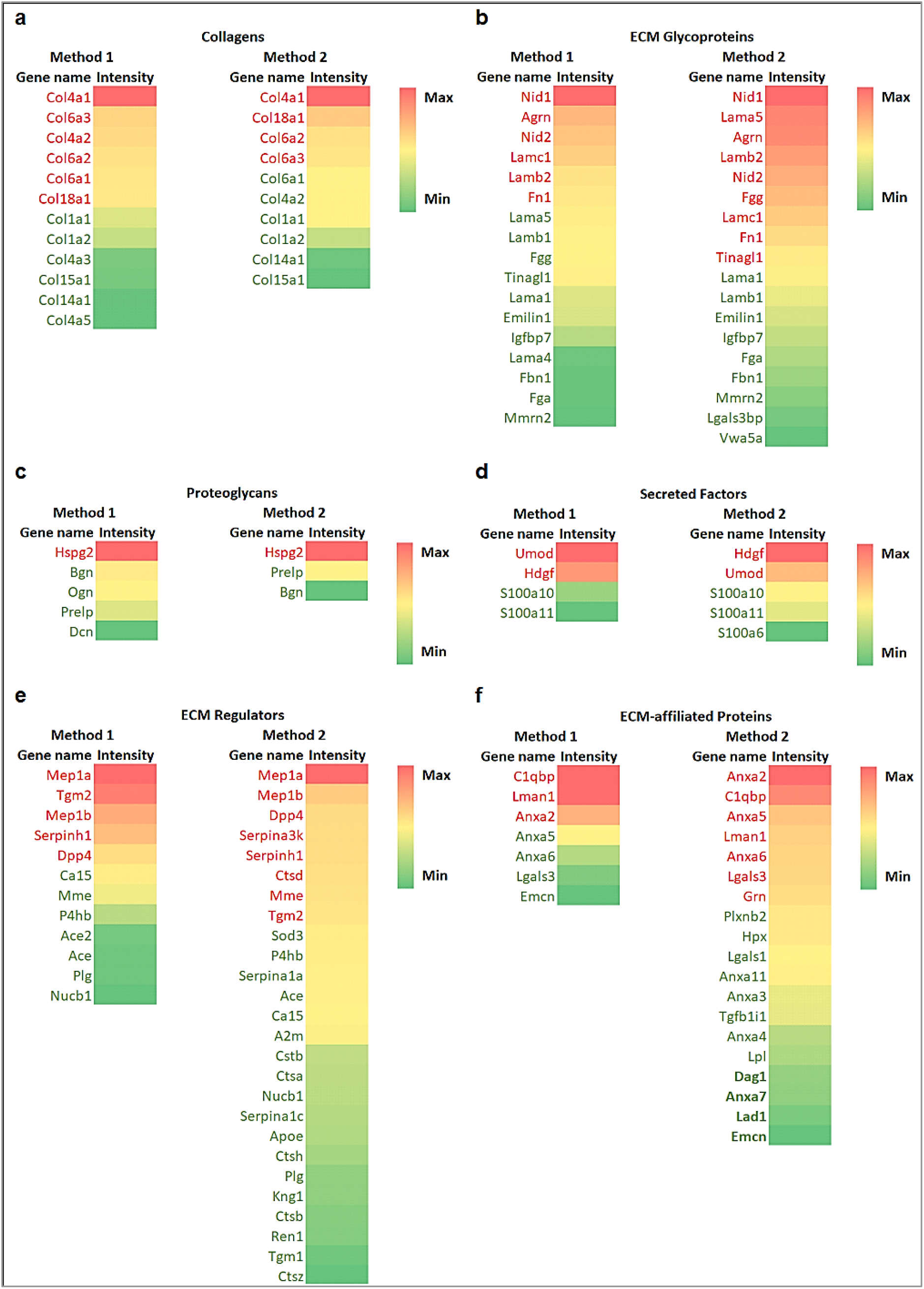
Heatmap of quantifiable proteins in Methods 1 and 2 comparing the abundance of each matrisome protein. Color coding in the heatmaps blocks depicts the variation between the maximum to minimum observed LQF intensity for each matrisome category and protein extraction method. The proteins’ names shown by red and green fonts are expressed above and below the average LFQ values in each matrisome category&extraction method, respectively.

The comparative analysis of the relative efficiency of Methods 1 and 2 in providing the samples for LFQ, revealed the following. 51 matrisome proteins (10 collagens, 16 ECM glycoprotein, 3 proteoglycans, 11 ECM regulator, 7 ECM affiliated and 4 secreted factors) were shared between the producst of both methods. Method 1 additionally allowed quantification of 6 matrisome proteins (2 collagens, 1 ECM glycoprotein, 2 proteoglycans and 1 ECM regulator). Method 2 added another 30 quantifiable matrisome proteins to the list (2 ECM glycoprotein, 15 ECM regulator and 12 ECM affiliated and 1 Secreted Factor). The comparative analysis of the quantification efficiency of the two studied protein extraction methods shows that there was no significant differences in abundances of shared matrisome proteins. However, a part of the quantifiable proteins was more abundant either in Method 1 or Method 2, while some were detected only in the specific method (**Table S4** in Supplementary Information). In general, Method 1 quantified more intensities of core matrisome proteins, but fewer ECM associated proteins compared to Method 2. For example, the abundance of some collagens (e.g Col1a2, Col6a1&2, Col18a1) were higher in Method 1, while Col4a3 and Col4a5 were only detected in Method 1. Similarly, some ECM glycoproteins (Lama1, Fgg, Fn1, & Igfbp7) and proteoglycans (Hspg2) were more abundant in Method 1. On the other hand, Method 2 revealed higher abundances of ECM-affiliated proteins (e.g. Anxa2, Anxa5 & Lgals3), ECM regulators (e.g. Ace, Dpp4, Meb1a & 1b) and secreted factors (Hdgf & S100a10 &11) while some of ECM-affiliated proteins (e.g. Anxa3, 4 &7, Hpx, Lad1), ECM regulators (e.g. Apoe, Serpina, Cts, Ren1, Kng1 and Sod3), secreted factors (S100a6) were only quantified by Method 2.

Furthermore, we have quantitatively compared the amount of protein extraction levels between Method 1 and the fractions obtained from Method 2 (i.e., 3 fractions, refer to Figure 1). The LFQ comparison revealed that 49, 27 and 52 matrisome proteins were quantitatively compared between Method 2-F1 *vs* Method 1, Method 2-F2 *vs* Method 1 and Method 2-F3 *vs* Method 1, respectively (**Figure S2,** and **Table S4**). Benjamin Hochberg FDR correction showed that Ace, Mep1b & Ctsa (ECM Regulators, Matrisome-associated) were significantly higher in Method 2-F1 compare to Method 1, whereas Method 1 obtained more abundances of core matrisome proteins including ECM Glycoproteins such as Nid1, Nid2, Tinagl1 & Agrn.

There were huge FC differences for some matrisome proteins in the comparison between Method 2-F2 and Method 1. Serpina1a & A2m are Matrisome-associated ECM Regulators and were 49.8 (*p*-value<0.0001) and 20.1 (*p*-value<0.001) folds higher in Method 2-F2, respectively. While Nid1 & Lamb1 are Core matrisome ECM Glycoproteins and were 64.4 (*p*-value<0.001) and 33.3 (*p*-value<0.001) folds higher in Method 1, respectively, compare to Method 2-F2.

## DISCUSSION

The current understanding of matrisome protein composition and how it is regulated during pathophysiological processes remaines very limited. One of the major obstacles is the optimization of matrisome protein extraction & characterization methods. Many studies, which examined kidney tissues of different species, used mainly decellularization approach that originates from tissue engineering and involves removal of cellular proteins followed by matrisome protein examination from the decellularized scaffolds ^55–57^. However, long detergent incubations in decellularization methods may cause degradation of some matrisome proteins ^58^.

The leading source of the information about ECM composition of human and mouse organs and tissues, the Matrisome Database, does not include the specification for mouse kidneys. Recently, the first reports on mouse renal matrisome have emerged, however, no quantitative data as well as no analysis of loosely bound ECM proteins were presented ^17,52^ Lack of such information impedes the progress in various types of kidney research, including the analysis of the pathological conditions that affect ECM metabolism.

Therefore, in current study, we aimed to explore mouse kidney matrisome using two widely used ECM enrichment methods and comprehensively compare the results using proteomics data analysis. To reach this goal, we used two extraction methods to enrich matrisome proteins. The first method was a commercially available Millipore Compartment Protein fractionation kit. This method biochemically separates subcellular proteins and enriches matrisome proteins in the insoluble pellet at the end step of the extraction series. The protocol for Method 1 is well documented in previous studies, and many ECM proteins were identified by using this extraction method. For example, Naba et al 2012^9^ identified 100 matrisome proteins in murine lung and colon, Schiller et al 2015^59^ identified 435 matrisome proteins in healthy mouse lungs and Gocheva et al 2017^16^ identified 113 matrisome proteins. Until recently, this method has not been used to identify matrisome proteins of murine kidneys. McCabe et al 2021^17^, and Lipp et al 2021^52^ identified 114 and 110 ECM proteins, respectively, using Millipore Compartment Fractionation.

In our study by using Millipore Compartment Fractionation (Method 1), we could detect 83 matrisome proteins of which 16 proteins were not previously presented in the Matrisome Database. It is important to note that, this method only results in one ECM-enriched fraction, while the loosely bound & soluble matrisome proteins may potentially remain in the cellular fractions. To improve the ECM protein isolation, we applied another widely used matrisome enrichment method, a sequential extraction approach (Method 2). This method allowed extraction of loosely bound or soluble matrisome proteins with high salt buffer before removal of cellular components by SDS and enriched matrisome fractions by solubilizing insoluble pellet with Gu-HCl ^60^. Similarly to our Method 2, Massey et al 2017 ^20^ identified 79 matrisome proteins from all three fractions of mouse liver. This extraction approach has shown to be a more comprehensive method to identify matrisome proteins but has not been optimized for kidney tissues. With this sequential approach, we have identified 105 matrisome proteins, of which, 22 proteins were yet to be listed in currently available MD (http://matrisomeproject.mit.edu/).

Meanwhile, we compared identified mouse matrisome proteins by studies of McCabe *et al*^17^, Lipp *et al*^52^ and Lui *et al*^49^, and identified human matrisome proteins by studies of Louzao-Martinez *et al*^53^ and Randles *et al*^54^. The comparsion provides differences between the lists of identified mouse and human specific matrisome proteins. In the combined list, we could identify 95 matrisome proteins only in mouse and 44 only in human kidneys. Interestingly, 5 of matrisome proteins shown in this study (**Table S2**) named Lgals3bp (glycoprotein), Mfge8 (glycoprotein) Emcn (ECM affiliated), Apoe (ECM regulator) and Sod3 (ECM regulator), were previously identified only in human kidneys^54^. However, among murine renal reports, only our study could identify these proteins in mouse kidneys. Hence, our study shows the necessity of using different approaches in identification of matrisome proteins to form a robust matrisome protein list.

To identify and quantify the extracted proteins from Methods 1 and 2, we performed a discovery proteomics study known as shotgun proteomics, which uses bottom-up approaches based on unbiased analysis. To compare the quantities of proteins in each method, we applied LFQ data generated from MaxQuan analysis ^25^. It is a widely used protein quantitation method in which the analysis can be performed either based on chromatographic ion intensities or based on spectral counting of identified proteins ^61^. It provides high throughput and does not have limitations of labelled based quantifications such as high complexity of sample preparation, requirement of high concentrations of samples and incomplete labelling^12,61^. In this study, we have successfully quantified 57 matrisome proteins in Method 1 and 81 in Method 2 using label-free based quantifications (LFQ). Some of the quantifiable proteins were more abundant either in Method 1 or Method 2. Our study showed that Method 1 could detect more intensities of core matrisome proteins, but fewer ECM associated proteins compared to Method 2.

To show the benefit of each extraction method which can be used as a guideline to study different matrisome proteins in different kidney diseases, we compared Method 2 fractions with Method 1, seperately. For example, IgA nephropathy (IgAN), which is one of the most prevalent chronic glomerular diseases, had significantly more matrisome proteins (Col4a1, Lamb1, Hspg2, Emilin, Fgg & Fbln) in diseased patients compared to healthy patients ^56^. Those proteins could be better detected using Method 1, while Col4a1 and Col15a1, which can be detected in both methods, were also elevated in IgAN, could be better detected in Method 1 or Method 2-F3. In renal fibrosis, the hallmark of CKDs, it is known that Col 1, 2, 3, 5, 6, 7 and 15, Fn1, Dcn and Bgn are accumulated in kidney ECM^23^. We could detect these matrisome proteins by both methods, while the abundances of some of them were higher in Method 1. The Method 2-F3 and Method 1 comparison also showed a majority of higher abundance proteins are Matrisome-associated (e.g., ECM-affiliated Proteins and ECM Regulators) whereas Core matrisome proteins (e.g., ECM Glycoproteins and Collagens) were higher in Method 1. Taken together, although Method 1 was powerful in extracting some of the matrisome proteins in higher abundances, Method 2 could quantify more matrisome proteins. Our observations also indicate that specific extraction methods may be required for the enrichment of specific matrisome proteins.

Although many matrisome proteins identified in our experiment have also been previously identified in other studies, we were also able to detect additional matrisome proteins. The most possible explanation of these extra findings is the use of different extraction methods. For example, LOX enzymes which are involved in collagen cross-linking, could not be detected in our extraction methods, but detected in another mouse kidney proteomic study where Millipore Compartment Fraction kit was used with combination of a modified deglycosylation/protein digestion processes ^17,52^. In addition, removal of glycosaminoglycans (GAGs) by digestion enzymes such as chondroitinase and heparinase, was suggested to improve peptide identification^60,62,63^. However, McCabe et al 2021^17^ reported that using GAG digesting enzymes only improved proteoglycan identification but not matrisome proteins. It is important to note that this study was performed on healthy kidney tissues and it is known that GAGs deposition in ECM is increased during the diseases, e.g. fibrosis^64,65^. Hence, the improvement of analysis by GAG digesting enzymes is still unclear, our study is limited with using only PNGaseF to remove glycans. We would suggest performing a pilot study using GAG digesting enzymes to analyse whether it improves the matrisome protein identification before applying it to all samples.

In conclusion, the better understanding of mouse kidney matrisome would provide more opportunities to perform studies in the areas of kidney disease modelling, kidney development biology, aging and tissue engineering. In this study, we successfully achieved a knowledge progress towards understanding the composition of mouse kidney matrisome and identifying proteins which were not listed as matrisome proteins. In addition, this study showed the importance of using different protein extraction steps to fractionate matrisome proteins for a better understanding of mouse kidney matrisome composition. For example, Method 2 which fractionated matrisome proteins as soluble and insoluble ECM proteins could identify and quantify more matrisome proteins in healthy murine kidneys. Method 1 which is based on obtaining one insoluble fraction, could not detect matrisome proteins as many as Method 2. Hence, we suggest to obtain different fractions of matriosome proteins to discover more ECM proteins for further studies of kidney ECM turnover in normal and pathological conditions, and also to idenfy novel drug targets and biomarkers for the treatment and diagnostics of chronic kidney diseases.

## EXPERIMENTAL PROCEDURES

### Animal preparation and tissue extraction

Murine kidney tissues were obtained via the post mortem animal tissues sharing program encouraged and approved by the UNSW Animal Care and Ethics Committee (ACEC). The WT C57BL/6 mice (female, aged 4–10 weeks, Australian BioResource) were housed in a stable environment at 21 ± 2 °C with a 12h/12h light-dark cycle. On the day of the experiment, the animals were anesthetized with 4% vaporized isoflurane delivered into an induction chamber and euthanized by cervical dislocation. First, the retinas were collected for the main experiment approved by the UNSW ACEC. After that, the animal bodies were placed on ice, underwent further dissection and kidneys’ extirpation. The obtained kidneys were kept on ice and one of the kidneys per animal were quickly transversely cut. Then, each half of a kidney (~50 mg) per animal was used for each protein extraction.

### Protein extraction

Two methods of protein extraction were employed in this study. For each method, three biological replicates were used (**Figure 1**).

#### Method 1

Millipore Compartment Protein Extraction Kit (Merck-Millipore, Cat. #2145) was used to deplete cytosolic, nuclear, membrane and cytoskeletal proteins and to enrich ECM proteins as described in Naba *et al*, 2015 ^66^. The obtained ECM enriched pellets were washed with 1x PBS (1.7 mM KH2PO4, 5 mM Na2HPO4, 150 mM NaCl, 25 mM EDTA, pH 7.4) containing 1:100 (v:v) of protease inhibitors (PI) (Halt Protease Inhibitor Cocktail, Thermo Scientific, Cat. #78429) three times to remove detergents and stored at −20 °C.

#### Method 2

Enrichment of ECM proteins was performed by the following protocol described in Barallobre-Baierro *et al*, 2017 ^60^. Briefly, half kidney samples were diced into 2-3mm pieces and washed five times with ice-cold 1x PBS containing 1:100 (v:v) of PI (Halt Protease Inhibitor Cocktail, Thermo Scientific, Cat. #78429) to minimize blood contamination. To extract ECM-associated, loosely bound ECM proteins and newly synthesized ECM proteins, washed samples were incubated in NaCl buffer (0.5 M NaCl, 10 mM Tris-HCl and 25 mM EDTA, pH 7.5 and 1:100 (v:v) of PI) for 1h at room temperature (RT) by vortexing at speed 65 rpm (Stuart Orbital Shaker). Samples were centrifuged at 16,000 x g for 10 min at 4 °C and supernatant was saved as Fraction 1. The pellet was treated with SDS buffer (0.1 % SDS, 25 mM EDTA and 1:100 (v:v) of PI) by vortexing at RT for 16 h at speed 65 rpm and was centrifuged at 16,000 x g for 10 min at 4 °C to separate intracellular proteins. Then, pellets were treated with GuHCl buffer (4 M guanidine hydrochloride, 50 mM Na acetate, 25 mM EDTA, pH 5.8, and 1:100 (v:v) of PI) by vortexing at RT for 72 h at speed 225 rpm to solubilise ECM proteins. Supernatants after centrifugation at 16,000 × g for 10 min at 4 °C were saved as Fraction 2 and pellets were saved as Fraction 3 after washing three times with ice-cold 1x PBS containing 1:100 (v:v) of PI. All saved fractions were stored at −20 °C.

#### Protein precipitation

Obtained pellet from Method 1 and three fractions from Method 2 (Figure 1b) were precipitated in EtOH separately to remove detergents/agents. 10 times volume of ice-cold 100% EtOH was added to each fraction and incubated overnight at −20 °C. Protein precipitates were obtained by centrifugation at 16,000 x g for 10 min at 4°C and after by drying pellets. Dried pellets were resuspended in the buffer of downstream process.

#### Deglycosylation and In-solution digestion

The method for deglycosylation and in-solution digestion was followed as described in Naba *et al*, 2015 ^66^. Briefly, obtained pellets for each fraction after protein precipitation were resuspended by adding the appropriate volume (50ul/5-10mg dry weight) of 8 M urea and DTT (Dithiotreitol) at a final concentration of 10mM. Samples were incubated with continuous agitation at 150 rpm (Stuart Orbital Shaker) for 2 hr at 37°C.

Alkylation was done by adding IAA (iodoacetamide) to a final concentration of 25 mM. To complete alkylation, the DTT: IAA ratio was adjusted as 1: 2.5 and incubated in the dark for 30 min at RT.

Deglycosylation was performed by diluting urea to 2M using 100 mM ammonium bicarbonate, pH 8.0 and adding 2ul/5-10mg dry weight (DW) of PNGaseF (Peptide-N-Glycosidase F) (New England Biolabs, Cat. #P0704S). Samples were incubated with continuous agitation at 150 rpm (Stuart Horizontal Shaker) for 2 hr at 37°C.

Digestion was continued with diluting samples with 100 mM ammonium bicarbonate pH 8.0 to obtain 1 M urea. Then, 2ul/5-10mg DW of Trypsin/Lys-C (Endoproteinase LysC) Mix (Promega, Cat #V5071) was added to samples and incubated with continuous agitation at 150 rpm for overnight at 37°C. The digestion reaction was inactivated with freshly prepared 8ul/5-10mg DW 50% TFA (Trifluoro-acetic acid) at a final concentration of 1% of TFA. The acidified samples were centrifuged at 16,000 x g for 5 min at RT and the supernatants were saved for desalting steps.

Desalting was performed by using Pierce C18 stage tips (Thermo Scientific, Cat, #SP301). Prior to proteomics analysis, desalted peptides were eluted with freshly prepared 60% acetonitrile, 0.1% TFA, followed by concentration in a vacuum concentrator. Samples then were resuspended in freshly prepared 3% acetonitrile, 0.1% TFA and analyzed in LC-MS/MS.

### LC-MS/MS and Data Analysis

Digested peptides were separated by nanoLC using an Ultimate nanoRSLC UPLC and autosampler system (Dionex, Amsterdam, Netherlands). Samples (2.5 μl) were concentrated and desalted onto a micro C18 precolumn (300 μm x 5 mm, Dionex) with H2O:CH3CN (98:2, 0.1 % TFA) at 15 μl/min. After a 4 min wash the pre-column was switched (Valco 10 port UPLC valve, Valco, Houston, TX) into line with a fritless nano column (75μ x ~20 cm) containing C18AQ media (1.9μ, 120 Å Dr Maisch, Ammerbuch-Entringen Germany) manufactured according to Gatlin. Peptides were eluted using a linear gradient of H_2_O:CH_3_CN (98:2, 0.1 % formic acid) to H_2_O:CH_3_CN (64:36, 0.1 % formic acid) at 200 nl/min over 30 min. High voltage 2000 V was applied to low volume Titanium union (Valco) with the column oven heated to 45°C (Sonation, Biberach, Germany) and the tip positioned ~ 0.5 cm from the heated capillary (T=300°C) of a QExactive Plus (Thermo Electron, Bremen, Germany) mass spectrometer. Positive ions were generated by electrospray and the QExactive operated in data-dependent acquisition mode (DDA).

A survey scan m/z 350-1750 was acquired (resolution = 70,000 at m/z 200, with an accumulation target value of 1,000,000 ions) and lock mass enabled (m/z 445.12003). Up to the 10 most abundant ions (>80,000 counts, underfill ratio 10%) with charge states > +2 and <+7 were sequentially isolated (width m/z 2.5) and fragmented by HCD (NCE = 30) with a AGC target of 100,000 ions (resolution = 17,500 at m/z 200). M/z ratios selected for MS/MS were dynamically excluded for 30 seconds.

MS raw files were analyzed by the MaxQuant software ^67^ (version 2.0.3.0), and peak lists were searched against the mouse UniProt FASTA database (version Aug 2019), and a common contaminants database by the Andromeda search engine ^68^. For protein identification and LFQ quantification, fractions of Method 2 were nominated as fractions in MaxQuant ^69^to have a merged output of fractions of the same sample. As fixed modification carbamidomethylation (C) and as variable modifications, oxidation (M,P), acetyl (protein N-term), hydroxyproline, deamidation (N,Q) were used. False discovery rate was set to 1% for proteins and peptides (minimum length of 7 amino acids) and was determined by searching a reverse database. Enzyme specificity was set as trypsin & lys-C, and a maximum of two missed cleavages were allowed in the database search. Peptide identification was performed with an allowed precursor mass deviation up to 4.5 ppm after time-dependent mass calibration and an allowed fragment mass deviation of 20 ppm. For LFQ, in MaxQuant, the minimum ratio count was set to two. For matching between runs, the retention time alignment window was set to 30 min and the match time window was 1 min.

For protein identification in Method 1 vs Method 2, in the first round of matrisome protein identification, unique peptides, which were presented at least in 2 of the biological replicates, were searched using “Matrisome Annotator” in the MD (http://matrisomeproject.mit.edu/). The matrisome proteins were classified as core matrisome proteins (collagens, ECM Glycoproteins & Proteoglycans) and ECM associated proteins (ECM-affiliated, ECM Regulators & Secreted Factors). In the second round, the protein list obtained from LC-MS/MS was matched with UniProt Database for *Mus musculus* and possible matrisome proteins were searched based on their subcellular locations using “extracellular matrix”, “extracellular space”, “basement membrane”, “secreted”, “lysosome” and “exosome”. To verify potential candidates, their possible interactions with matrisome proteins were searched in protein interaction databases (BioGRID, STRING). In the final stage, the short list of proteins was searched in the literature to confirm that proteins have been located in ECM by other studies.

For protein quantification, LFQ intensity values were taken from the MaxQuant protein “Groups” table, which represented the values after inter-experiment normalization ^70^. Differential abundant analysis was performed using LFQ-Analyst online software (https://bioinformatics.erc.monash.edu/apps/LFQ-Analyst/)^25^. Matrisome proteins which had LFQ intensities in at least 2 of 3 biological replicates, were included in the comparison of Method 1 and Method 2 or Method 2-fractions. In LFQ-Analyst, parameters were set as Perseus-type imputation, adjusted p-value cutoff<0.05 and Log2 fold change (FC) cutoff=1 (i.e., 2-FC). The comparison was presented with FC and p-values. All p-values were corrected for multiple hypothesis testing using the Benjamini–Hochberg method. Proteins with p<0.05 and 2-FC increase or decease were presented as significantly different. The error bars correspond to the standard error of the mean.

LFQ analysis to compare fractions of Method 2 and Method 1 were performed separately for each fraction, and same parameters, which were used to compare Method 1 & 2, were used but fractions were set as “separate method” instead of setting as “fractions”.

## ABBREVIATIONS

ECM: Extracellular matrix
matrisome: ECM proteome
LC-MS/MS: liquid chromatographymass spectrometry
MD: Matrisome database
CKDs: chronic kidney diseases
GPs: glycoproteins
PGs: proteoglycans
SDS: sodium dodecyl sulphate
LFQ: label free quantification.

## ACKNOWLEDGEMENT

Authors thank Prof Ewa Goldys, Prof Carol Pollock, A/Prof Sonia Saad, Dr Wolfgang Jarolimek, Dr Long Nguyen and Dr Yimin Yao for helpful discussions. Mass spectrometric results were obtained at the Bioanalytical Mass Spectrometry Facility within the Mark Wainwright Analytical Centre of the University of New South Wales. We thank Dr Ling Zhong for the support in mass spectrometric analysis. We are grateful to Dr Tianruo Guo and Ms Madhuvanthi Muralidharan (UNSW) for sharing of the animal tissues and Ms Naomi Craig for the professional support in organizing the tissue sharing workflow. This work was supported by the funding from NSW Health Department, UNSW and Pharmaxis (NSW Health PhD Partnership Program Scholarship to U.R. and A.G.).

## AUTHORS CONTRIBUTIONS

U.R. and A.G. were responsible for hypothesis development, design or the experiments, and final data integrity and interpretation. U.R., S.B.A., and S.A. performed all proteomic analyses. All authors contributed to the conceptual design of the manuscript and to data interpretation. U.R. wrote the manuscript, and all authors edited and approved the final manuscript submission.

## DATA AVAILABILITY STATEMENT

All relevant data generated and (or) analyzed in the current study is available in the manuscript and Supproting Information.

