## Supplementary Information for "IDENTIFICATION AND QUANTIFICATION OF MATRISOME PROTEINS OF MOUSE KIDNEYS"

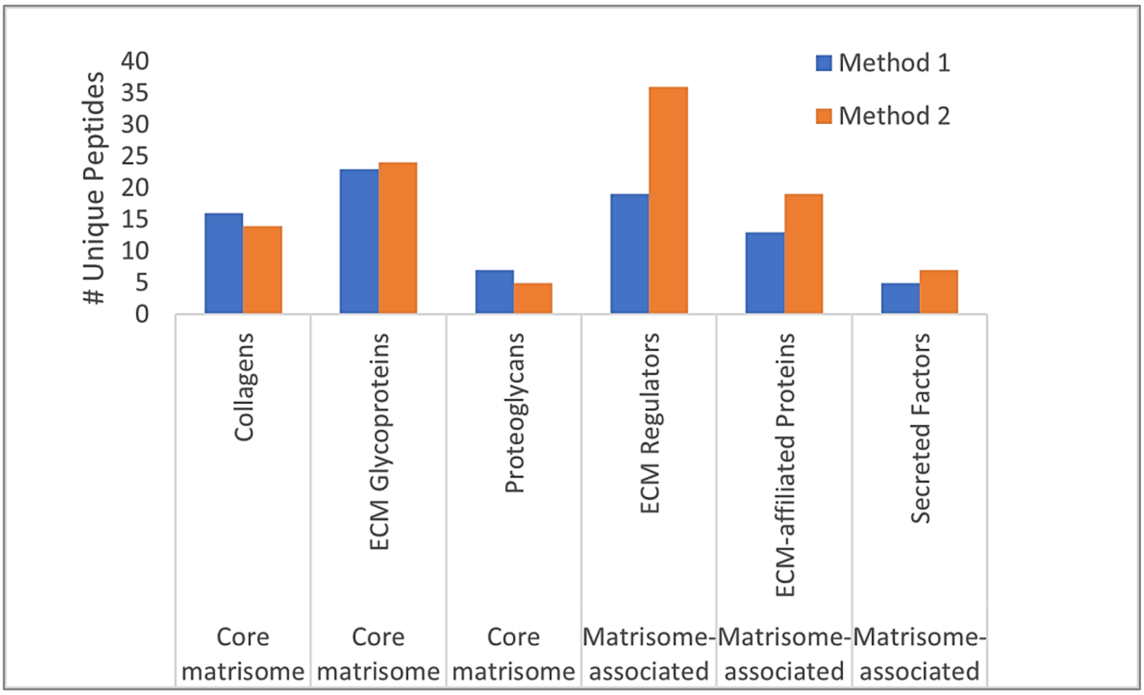


**Figure S1**. Unique peptides identifying protein IDs in Method 1 and Method 2. It compares number of unique peptides in each category and division in Method 1 and Method 2. Matrisome proteins were classified as Collagens, ECM Glycoproteins, Proteoglycans, ECM-affiliated, ECM Regulators and Secreted Factors.

**Table S1.** The list of matrisome proteins identified in the current study in at least two of the biological replicates. Rows highlighted in yellow describe identified matrisome proteins which were not previously listed in MD. Abbreviations indicate the following: Divisions of matrisome: CM – core matrisome proteins; MAP – matrisome-associated proteins. Categories of matrisome: Col – collagens, GP – ECM Glycoproteins, PG – Proteoglycans, ECMR – ECM Regulators, ECMAP – ECM-affiliated proteins, SF – secreted factors. Ref- references: MD- the Matrisome Database; numbers are the literature references.

| Protein ID | Division | Category | Protein Name | Gene Symbol | Identification by Methods 1 (M1) and 2 (M2) | | Ref |
|---|---|---|---|---|---|---|---|
| | | | | | M1 | M2 | |
| P11087 | CM | Col | Collagen alpha-1(I) chain | Col1a1 | + | + | MD |
| Q01149 | CM | Col | Collagen alpha-2(I) chain | Col1a2 | + | + | MD |
| P08121 | CM | Col | Collagen alpha-1(III) chain | Col3a1 | + | + | MD |
| P02463 | CM | Col | Collagen alpha-1(IV) chain | Col4a1 | + | + | MD |
| P08122 | CM | Col | Collagen alpha-2(IV) chain | Col4a2 | + | + | MD |
| Q9QZS0 | CM | Col | Collagen alpha-3(IV) chain | Col4a3 | + | - | MD |
| Q9QZR9 | CM | Col | Collagen alpha-4(IV) chain | Col4a4 | + | - | MD |
| Q9CPW5 | CM | Col | Collagen alpha-5(IV) chain | Col4a5 | + | + | MD |
| O88207 | CM | Col | Collagen alpha-1(V) chain | Col5a1 | + | + | MD |
| Q3U962 | CM | Col | Collagen alpha-2(V) chain | Col5a2 | + | + | MD |
| Q04857 | CM | Col | Collagen alpha-1(VI) chain | Col6a1 | + | + | MD |
| Q02788 | CM | Col | Collagen alpha-2(VI) chain | Col6a2 | + | + | MD |
| Q61001 | CM | Col | Collagen alpha-3(VI) chain | Col6a3 | + | + | MD |
| Q80X19 | CM | Col | Collagen alpha-1(XIV) chain | Col14a1 | + | + | MD |
| O35206 | CM | Col | Collagen alpha-1(XV) chain | Col15a1 | + | + | MD |
| P39061 | CM | Col | Collagen alpha-1(XVIII) chain | Col18a1 | + | + | MD |
| A2ASQ1 | CM | GP | Agrin | Agrn | + | + | MD |
| Q9QZZ6 | CM | GP | Dermatopontin | Dpt | + | - | MD |
| Q91VF5 | CM | GP | EMI domain-containing protein 1 | Emid1 | + | + | MD |
| Q99K41 | CM | GP | EMILIN-1 | Emilin1 | + | + | MD |
| Q61554 | CM | GP | Fibrillin-1 | Fbn1 | + | - | MD |
| Q61555 | CM | GP | Fibrillin-2 | Fbn2 | - | + | MD |
| E9PV24 | CM | GP | Fibrinogen alpha chain | Fga | + | + | MD |
| Q8VCM7 | CM | GP | Fibrinogen gamma chain | Fgg | + | + | MD |
| P11276 | CM | GP | Fibronectin | Fn1 | + | + | MD |
| A2ASQ1 | CM | GP | Progranulin | Grn | - | + | MD |
| Q61581 | CM | GP | Insulin-like growth factor-binding protein 7 | Igfbp7 | + | + | MD |
| P19137 | CM | GP | Laminin subunit alpha-1 | Lama1 | + | + | MD |
| P97927 | CM | GP | Laminin subunit alpha-4 | Lama4 | + | + | MD |
| Q61001 | CM | GP | Laminin subunit alpha-5 | Lama5 | + | + | MD |
| P02469 | CM | GP | Laminin subunit beta-1 | Lamb1 | + | + | MD |
| Q61292 | CM | GP | Laminin subunit beta-2 | Lamb2 | + | + | MD |
| P02468 | CM | GP | Laminin subunit gamma-1 | Lamc1 | + | + | MD |
| Q07797 | CM | GP | Galectin-3-binding protein | Lgals3bp | + | + | 26 |
| Q8K4G1 | CM | GP | Latent TGF-b-binding protein 4 | Ltbp4 | - | + | MD |
| P21956 | CM | GP | Lactadherin | Mfge8 | + | + | MD |
| A6H6E2 | CM | GP | Multimerin-2 | Mmrn2 | + | + | MD |
| P10493 | CM | GP | Nidogen-1 | Nid1 | + | + | MD |
| O88322 | CM | GP | Nidogen-2 | Nid2 | + | + | MD |
| Q91V88 | CM | GP | Nephronectin | Npnt | - | + | MD |
| Q99JR5 | CM | GP | Tubulointerstitial nephritis antigen-like | Tinagl1 | + | + | MD |
| P29788 | CM | GP | Vitronectin | Vtn | + | - | MD |
| Q8R2Z5 | CM | GP | von Willebrand factor A domain-containing protein 1 | Vwa1 | + | - | MD |
| Q99KC8 | CM | GP | von Willebrand factor A domain-containing protein 5A | Vwa5a | - | + | MD |
| Q99MQ4 | CM | PG | Asporin | Aspn | + | - | MD |
| P28653 | CM | PG | Biglycan | Bgn | + | + | MD |
| P28654 | CM | PG | Decorin | Dcn | + | + | MD |
| Q05793 | CM | PG | Heparan sulfate proteoglycan core protein | Hspg2 | + | + | MD |
| P51885 | CM | PG | Lumican | Lum | + | + | MD |
| Q62000 | CM | PG | Mimecan | Ogn | + | - | MD |
| Q9JK53 | CM | PG | Prolargin | Prelp | + | + | MD |
| Q8JZZ0 | MAP | PG | Alpha-2-macroglobulin; Alpha-2-macroglobulin 165 kDa subunit; Alpha-2-macroglobulin 35 kDa subunit | A2m | - | + | MD |
| P09470 | MAP | ECMR | Angiotensin-converting enzyme | Ace | + | + | 27 |

| | | | | | | | |
|---|---|---|---|---|---|---|---|
| Q8R0I0 | MAP | ECMR | Angiotensin-converting enzyme 2 | Ace2 | + | + | 28 |
| O35598 | MAP | ECMR | Disintegrin and metalloproteinase domain-containing protein 10 | Adam10 | + | + | MD |
| P08226 | MAP | ECMR | Apolipoprotein E | Apoe | - | + | 29 |
| P50429 | MAP | ECMR | Arylsulfatase B | Arsb | - | + | 30 |
| Q99N23 | MAP | ECMR | Carbonic anhydrase 15 | Ca15 | + | + | 31 |
| Q62426 | MAP | ECMR | Cystatin-B | Cstb | - | + | MD |
| P16675 | MAP | ECMR | Lysosomal protective protein | Ctsa | + | + | MD |
| P10605 | MAP | ECMR | Cathepsin B | Ctsb | + | + | MD |
| P18242 | MAP | ECMR | Cathepsin D | Ctsd | + | + | MD |
| P49935 | MAP | ECMR | Pro-cathepsin H | Ctsh | + | + | MD |
| Q9WUU7 | MAP | ECMR | Cathepsin Z | Ctsz | - | + | MD |
| P28843 | MAP | ECMR | Dipeptidyl peptidase 4 | Dpp4 | + | + | 32 |
| P19221 | MAP | ECMR | Prothrombin | F2 | - | + | MD |
| Q571E4 | MAP | ECMR | N-acetylgalactosamine-6-sulfatase | Galns | - | + | 33 |
| Q8BFR4 | MAP | ECMR | N-acetylglucosamine-6-sulfatase | Gns | - | + | 34 |
| Q9ESB3 | MAP | ECMR | Histidine-rich glycoprotein | Hrg | - | + | MD |
| Q61703 | MAP | ECMR | Inter-alpha-trypsin inhibitor heavy chain H2 | Itih2 | - | + | MD |
| O08677 | MAP | ECMR | Kininogen-1 | Kng1 | - | + | MD |
| P28825 | MAP | ECMR | Meprin A subunit alpha | Mep1a | + | + | MD |
| Q61847 | MAP | ECMR | Meprin A subunit beta | Mep1b | + | + | MD |
| Q61391 | MAP | ECMR | Neprilysin | Mme | + | + | 35 |
| Q02819 | MAP | ECMR | Nucleobindin-1 | Nucb1 | + | + | 36 |
| P09103 | MAP | ECMR | Protein disulfide-isomerase | P4hb | + | + | 37 |
| P20918 | MAP | ECMR | Plasminogen | Plg | + | + | MD |
| P06281 | MAP | ECMR | Renin-1 | Ren1 | - | + | 38 |
| P07758 | MAP | ECMR | Alpha-1-antitrypsin 1-1 | Serpina1a | - | + | MD |
| P22599 | MAP | ECMR | Alpha-1-antitrypsin 1-2 | Serpina1b | - | + | MD |
| Q00896 | MAP | ECMR | Alpha-1-antitrypsin 1-3 | Serpina1c | - | + | MD |
| P07759 | MAP | ECMR | Serine protease inhibitor A3K | Serpina3k | - | + | MD |
| Q60854 | MAP | ECMR | Serpin B6 | Serpinb6 | - | + | 39 |
| P19324 | MAP | ECMR | Serpin H1 | Serpinh1 | + | + | MD |
| O09164 | MAP | ECMR | Extracellular superoxide dismutase | Sod3 | + | + | 40 |
| Q9JLF6 | MAP | ECMR | Protein-glutamine gamma-glutamyltransferase K | Tgm1 | + | + | MD |
| P21981 | MAP | ECMR | Protein-glutamine gamma-glutamyltransferase 2 | Tgm2 | + | + | MD |
| P10107 | MAP | ECMAP | Annexin A1 | Anxa1 | - | + | MD |
| P07356 | MAP | ECMAP | Annexin A2 | Anxa2 | + | + | MD |
| O35639 | MAP | ECMAP | Annexin A3 | Anxa3 | - | + | MD |
| P97429 | MAP | ECMAP | Annexin A4 | Anxa4 | - | + | MD |
| P48036 | MAP | ECMAP | Annexin A5 | Anxa5 | + | + | MD |
| P14824 | MAP | ECMAP | Annexin A6 | Anxa6 | + | + | MD |
| Q07076 | MAP | ECMAP | Annexin A7 | Anxa7 | - | + | MD |
| P97384 | MAP | ECMAP | Annexin A11 | Anxa11 | - | + | MD |
| O35658 | MAP | ECMAP | Complement component 1 Q subcomponent-binding protein | C1qbp | + | + | 41,42 |
| Q62165 | MAP | ECMAP | Dystroglycan | Dag1 | + | + | 43 |
| Q9R0H2 | MAP | ECMAP | Endomucin | Emcn | + | + | MD |
| Q91X72 | MAP | ECMAP | Hemopexin | Hpx | - | + | MD |
| P57016 | MAP | ECMAP | Ladinin-1 | Lad1 | + | + | 44 |
| P16045 | MAP | ECMAP | Galectin-1 | Lgals1 | + | + | MD |
| P16110 | MAP | ECMAP | Galectin-3 | Lgals3 | + | + | MD |
| Q9D0F3 | MAP | ECMAP | Protein ERGIC-53 | Lman1 | + | + | MD |
| P11152 | MAP | ECMAP | Lipoprotein lipase | Lpl | + | + | 45 |
| B2RXS4 | MAP | ECMAP | Plexin-B2 | Plxnb2 | + | + | MD |

| | | | | | | | |
|---|---|---|---|---|---|---|---|
| Q62219 | MAP | ECMAP | TGF-b-1-induced transcript 1 protein | Tgfb1i1 | + | + | 46 |
| P01132 | MAP | SF | Pro-epidermal growth factor | Egf | - | + | MD |
| Q8CJ70 | MAP | SF | Interleukin-19 | Il19 | + | + | MD |
| P14069 | MAP | SF | Protein S100-A6 | S100a6 | - | + | MD |
| P08207 | MAP | SF | Protein S100-A10 | S100a10 | + | + | MD |
| P50543 | MAP | SF | Protein S100-A11 | S100a11 | + | + | MD |
| Q91X17 | MAP | SF | Uromodulin | Umod | + | + | 47 |
| P51859 | MAP | SF | Hepatoma-derived growth factor | Hdgf | + | + | 48 |

**Table S2.** A combined list of matrisome proteins in mouse and human kidneys. The mouse matrisome proteins were obtained from our data and from references [17,49,52] and human matrisome proteins were obtained from the studies [53,54]. Highlighted "X" refers to the protein identified in the corresponding study.

| | | Our data | McCabe 2021 | Lipp 2021 | Lui 2020 | Louzao-Martinez 2019 | Randles 2021 |
|---|---|---|---|---|---|---|---|
| **protein name** | **category** | **mouse kidney** | **mouse kidney** | **mouse kidney** | **mouse kidney** | **human kidney** | **human kidney** |
| COL1A1 | Collagens | X | X | X | X | X | X |
| COL1A2 | Collagens | X | X | X | X | X | X |
| COL2A1 | Collagens | | X | | | | |
| COL3A1 | Collagens | X | X | X | | X | X |
| COL4A1 | Collagens | X | X | X | X | X | X |
| COL4A2 | Collagens | X | X | X | X | X | X |
| COL4A3 | Collagens | X | X | X | X | | X |
| COL4A4 | Collagens | X | X | X | X | | X |
| COL4A5 | Collagens | | | X | X | | X |
| COL4A6 | Collagens | | | X | | | X |
| COL5A1 | Collagens | X | X | X | | | X |
| COL5A2 | Collagens | X | X | X | | | X |
| COL6A1 | Collagens | X | X | X | X | X | X |
| COL6A2 | Collagens | X | X | X | X | X | X |
| COL6A3 | Collagens | | | X | X | X | X |
| COL6A5 | Collagens | | X | | X | | |
| COL6A6 | Collagens | | X | | X | | |
| COL7A1 | Collagens | | | | | | X |
| COL8A1 | Collagens | | | | | | X |
| COL11A1 | Collagens | | X | | | | |
| COL11A2 | Collagens | | X | | | | |
| COL12A1 | Collagens | | X | X | X | X | X |
| COL14A1 | Collagens | X | X | X | X | X | X |
| COL15A1 | Collagens | X | X | X | X | X | X |
| COL16A1 | Collagens | | X | | | | X |
| COL18A1 | Collagens | X | X | X | X | X | X |
| COL24A1 | Collagens | | X | | | | |
| ADIPOQ | ECM Glycoproteins | | | | X | | |
| AEBP1 | ECM Glycoproteins | | X | | | | |
| AGRN | ECM Glycoproteins | X | X | X | X | X | X |
| AHSG | ECM Glycoproteins | | | | | | X |
| APOH | ECM Glycoproteins | | | | | | X |
| CRELD1 | ECM Glycoproteins | | | | | | X |
| CRELD2 | ECM Glycoproteins | | | | X | | |
| DPT | ECM Glycoproteins | X | X | X | | X | X |
| ECM1 | ECM Glycoproteins | | X | X | X | | X |
| EFEMP1 | ECM Glycoproteins | | X | | | | X |
| EFEMP2 | ECM Glycoproteins | | X | | | | |
| ELN | ECM Glycoproteins | | X | X | | X | X |
| EMID1 | ECM Glycoproteins | X | X | | | | |
| EMILIN1 | ECM Glycoproteins | X | X | X | X | X | X |
| FBLN1 | ECM Glycoproteins | | X | | X | X | X |
| FBLN2 | ECM Glycoproteins | | X | | | | |
| FBLN5 | ECM Glycoproteins | X | X | X | | X | X |
| FBN1 | ECM Glycoproteins | X | X | X | X | X | X |
| FBN2 | ECM Glycoproteins | X | | | | X | |
| FGA | ECM Glycoproteins | X | X | X | X | X | X |
| FGB | ECM Glycoproteins | | X | X | X | X | X |
| FGG | ECM Glycoproteins | X | X | X | X | X | X |
| FGL1 | ECM Glycoproteins | | | | | | X |
| FGL2 | ECM Glycoproteins | | | | | | X |
| FN1 | ECM Glycoproteins | X | X | X | X | X | X |
| FRAS1 | ECM Glycoproteins | | | X | X | X | X |
| IGFBP7 | ECM Glycoproteins | X | X | X | X | X | X |
| KCP | ECM Glycoproteins | | X | X | X | | X |
| LAMA1 | ECM Glycoproteins | X | X | X | X | X | X |
| LAMA2 | ECM Glycoproteins | | X | X | X | | X |
| LAMA3 | ECM Glycoproteins | X | X | X | X | | X |
| LAMA4 | ECM Glycoproteins | X | X | X | X | X | X |

| | | | | | | | |
|---|---|---|---|---|---|---|---|
| LAMA5 | ECM Glycoproteins | X | X | X | X | X | X |
| LAMB1 | ECM Glycoproteins | X | X | X | X | X | X |
| LAMB2 | ECM Glycoproteins | X | X | X | X | X | X |
| LAMB3 | ECM Glycoproteins | | X | | | | |
| LAMC1 | ECM Glycoproteins | X | X | X | X | X | X |
| LAMC3 | ECM Glycoproteins | | X | | X | | |
| LGALS3BP | ECM Glycoproteins | X | | | | | X |
| LTBP1 | ECM Glycoproteins | | X | X | | | |
| LTBP4 | ECM Glycoproteins | X | X | | X | | X |
| MATN2 | ECM Glycoproteins | | X | X | X | | X |
| MFAP2 | ECM Glycoproteins | | X | X | | X | |
| MFAP4 | ECM Glycoproteins | | X | | X | | X |
| MFAP5 | ECM Glycoproteins | | X | | X | X | |
| MFGE8 | ECM Glycoproteins | X | | | | | X |
| MMRN2 | ECM Glycoproteins | X | X | X | X | | X |
| NID1 | ECM Glycoproteins | X | X | X | X | X | X |
| NID2 | ECM Glycoproteins | X | X | X | X | X | X |
| NPNT | ECM Glycoproteins | X | X | X | X | X | |
| NTN1 | ECM Glycoproteins | | | | X | | X |
| NTN4 | ECM Glycoproteins | | X | | X | | |
| PAPLN | ECM Glycoproteins | | X | | X | | X |
| POSTN | ECM Glycoproteins | X | X | X | X | X | X |
| PXDN | ECM Glycoproteins | | X | X | X | | X |
| SBSPON | ECM Glycoproteins | | X | | | | X |
| SPARC | ECM Glycoproteins | | X | | | | X |
| TGFBI | ECM Glycoproteins | | X | X | X | X | X |
| THBS1 | ECM Glycoproteins | | X | | | X | X |
| THSD4 | ECM Glycoproteins | | X | | | | X |
| TINAG | ECM Glycoproteins | | | X | | X | X |
| TINAGL1 | ECM Glycoproteins | X | X | X | X | X | X |
| TNC | ECM Glycoproteins | | X | X | | X | X |
| TNXB | ECM Glycoproteins | | | X | | | |
| VTN | ECM Glycoproteins | X | X | X | X | X | X |
| VWA1 | ECM Glycoproteins | X | X | X | | | X |
| VWA2 | ECM Glycoproteins | | X | | X | | |
| VWA5A | ECM Glycoproteins | X | X | | X | | |
| VWF | ECM Glycoproteins | | X | | | X | X |
| A2M | ECM Regulators | X | X | | | X | X |
| ACE | ECM Regulators | X | | | | | |
| ACE2 | ECM Regulators | X | | | | | |
| ADAM8 | ECM Regulators | | | | X | | |
| ADAM9 | ECM Regulators | | | | | | X |
| ADAM10 | ECM Regulators | X | X | | X | | |
| ADAM17 | ECM Regulators | | | | X | | |
| ADAMTSL1 | ECM Regulators | | X | | | | |
| ADAMTSL4 | ECM Regulators | | X | | | | |
| ADAMTSL5 | ECM Regulators | | | | | | X |
| AGT | ECM Regulators | | X | | | | |
| AMBP | ECM Regulators | | X | | X | X | X |
| APOE | ECM Regulators | X | | | | | X |
| ARSB | ECM Regulators | X | | | | | |
| CA15 | ECM Regulators | X | | | | | |
| CASP14 | ECM Regulators | | | | | | X |
| CPN2 | ECM Regulators | | X | | X | | |
| CST3 | ECM Regulators | | X | | | | |
| CSTB | ECM Regulators | X | X | | X | | |
| CTSA | ECM Regulators | X | X | | X | | X |
| CTSB | ECM Regulators | X | X | | X | | X |
| CTSC | ECM Regulators | | X | | | | X |
| CTSD | ECM Regulators | X | X | | X | X | X |
| CTSF | ECM Regulators | | X | | | | |
| CTSH | ECM Regulators | X | X | | X | | X |
| CTSL | ECM Regulators | | X | | X | | |
| CTSS | ECM Regulators | | | | X | | X |
| CTSZ | ECM Regulators | X | X | | X | | X |
| DPP4 | ECM Regulators | X | | | | | |

| | | | | | | | |
|---|---|---|---|---|---|---|---|
| ELANE | ECM Regulators | | | | | | X |
| F13A1 | ECM Regulators | | X | | | | |
| F2 | ECM Regulators | X | X | | X | | |
| F3 | ECM Regulators | | | | | | X |
| F9 | ECM Regulators | | X | | X | | X |
| FAM20B | ECM Regulators | X | X | | X | | |
| GALNS | ECM Regulators | X | | | | | |
| GANAB | ECM Regulators | | | | | | X |
| GNS | ECM Regulators | X | | | | | |
| HRG | ECM Regulators | X | X | | X | X | X |
| HTRA1 | ECM Regulators | | X | | | | X |
| ITIH1 | ECM Regulators | | X | X | X | X | X |
| ITIH2 | ECM Regulators | X | X | | X | X | |
| ITIH4 | ECM Regulators | | X | | X | | X |
| ITIH5 | ECM Regulators | | X | | | | X |
| KNG1 | ECM Regulators | X | X | | X | | X |
| LOX | ECM Regulators | | X | | | | |
| LOXL1 | ECM Regulators | | X | | | | |
| LOXL2 | ECM Regulators | | | X | | | |
| LOXL4 | ECM Regulators | | X | | | | |
| MEP1A | ECM Regulators | X | X | X | X | | |
| MEP1B | ECM Regulators | X | X | X | X | | |
| MME | ECM Regulators | X | | | | | |
| MMP9 | ECM Regulators | | | | | | X |
| MUG2 | ECM Regulators | | X | | X | | |
| NGLY1 | ECM Regulators | | | | X | | |
| NUCB1 | ECM Regulators | X | | | | | |
| OGFOD2 | ECM Regulators | | | | X | | |
| P4HA1 | ECM Regulators | | X | | | | X |
| P4HB | ECM Regulators | X | | | | | |
| PLAU | ECM Regulators | | X | | | | |
| PLG | ECM Regulators | X | X | X | X | | X |
| PLOD1 | ECM Regulators | | | | | | X |
| PLOD2 | ECM Regulators | | | | | | X |
| PLOD3 | ECM Regulators | | | | | | X |
| PLSCR1 | ECM Regulators | | | | | | X |
| PRDX1 | ECM Regulators | | | | | | X |
| PRTN3 | ECM Regulators | | | | | | X |
| REN1 | ECM Regulators | X | | | | | |
| SERPINA1 | ECM Regulators | | | | | X | |
| SERPINA1A | ECM Regulators | X | X | X | X | | |
| SERPINA1B | ECM Regulators | X | X | | X | | |
| SERPINA1C | ECM Regulators | X | X | | X | | |
| SERPINA1D | ECM Regulators | | X | | X | | |
| SERPINA1E | ECM Regulators | | X | | X | | |
| SERPINA3 | ECM Regulators | | | | | X | |
| SERPINA3G | ECM Regulators | | | | X | | |
| SERPINA3K | ECM Regulators | X | X | | X | | |
| SERPINA3M | ECM Regulators | | X | | X | | |
| SERPINA3N | ECM Regulators | | X | | | | |
| SERPINA5 | ECM Regulators | | | | | | X |
| SERPINA6 | ECM Regulators | | X | | X | | |
| SERPINB1A | ECM Regulators | | X | | X | | |
| SERPINB6 | ECM Regulators | X | | | X | | |
| SERPINB12 | ECM Regulators | | | | | | X |
| SERPINC1 | ECM Regulators | | X | | X | X | X |
| SERPIND1 | ECM Regulators | | X | | X | | X |
| SERPINF1 | ECM Regulators | | X | | | | X |
| SERPINF2 | ECM Regulators | | X | | X | | X |
| SERPING1 | ECM Regulators | | X | | X | | X |
| SERPINH1 | ECM Regulators | X | X | X | X | | X |
| SOD1 | ECM Regulators | | | | | | X |
| SOD3 | ECM Regulators | X | | | | | X |
| ST14 | ECM Regulators | | X | | | | |
| TGM1 | ECM Regulators | X | X | X | X | | X |
| TGM2 | ECM Regulators | X | X | X | X | X | X |

| | | | | | | | |
|---|---|---|---|---|---|---|---|
| TGM3 | ECM Regulators | | | | | | X |
| TIMP3 | ECM Regulators | | X | | X | X | X |
| ANXA1 | ECM-affiliated Proteins | X | X | | X | X | X |
| ANXA2 | ECM-affiliated Proteins | X | X | X | X | | X |
| ANXA3 | ECM-affiliated Proteins | X | X | | X | | |
| ANXA4 | ECM-affiliated Proteins | X | X | | X | X | X |
| ANXA5 | ECM-affiliated Proteins | X | X | | X | X | X |
| ANXA6 | ECM-affiliated Proteins | X | X | X | X | X | X |
| ANXA7 | ECM-affiliated Proteins | X | X | X | X | X | X |
| ANXA9 | ECM-affiliated Proteins | | X | | | | |
| ANXA11 | ECM-affiliated Proteins | X | X | X | X | X | X |
| ANXA13 | ECM-affiliated Proteins | | | | X | | X |
| APCS | ECM-affiliated Proteins | | | | | | X |
| C1QBP | ECM-affiliated Proteins | X | | | | | |
| CCT2 | ECM-affiliated Proteins | | | | | | X |
| CCT6A | ECM-affiliated Proteins | | | | | | X |
| CD109 | ECM-affiliated Proteins | | | | | | X |
| CSPG4 | ECM-affiliated Proteins | | X | | X | | X |
| CXCL14 | ECM-affiliated Proteins | | | | | | X |
| DAG1 | ECM-affiliated Proteins | X | | | | | |
| EMCN | ECM-affiliated Proteins | X | | | | | X |
| FREM1 | ECM-affiliated Proteins | | | | | | X |
| FREM2 | ECM-affiliated Proteins | | X | X | X | X | |
| GPC4 | ECM-affiliated Proteins | | X | | X | | |
| GRN | ECM-affiliated Proteins | X | | | | | |
| HPX | ECM-affiliated Proteins | X | X | | X | | X |
| LAD | ECM-affiliated Proteins | X | | | | | |
| LGALS1 | ECM-affiliated Proteins | X | X | | X | X | X |
| LGALS2 | ECM-affiliated Proteins | | | | | | X |
| LGALS3 | ECM-affiliated Proteins | X | X | | | | X |
| LGALS8 | ECM-affiliated Proteins | | X | | | | X |
| LGALS9 | ECM-affiliated Proteins | | X | | X | | |
| LMAN1 | ECM-affiliated Proteins | X | X | X | X | X | X |
| LPL | ECM-affiliated Proteins | X | | | | | |
| MBL1 | ECM-affiliated Proteins | | X | | | | |
| MBL2 | ECM-affiliated Proteins | | X | | | | |
| MGP | ECM-affiliated Proteins | | | | | | X |
| MUC1 | ECM-affiliated Proteins | | | | | | X |
| PLXNB1 | ECM-affiliated Proteins | | | | | | X |
| PLXNB2 | ECM-affiliated Proteins | X | X | | X | | X |
| PLXND1 | ECM-affiliated Proteins | | | | X | | X |
| SDC4 | ECM-affiliated Proteins | | | | X | | |
| SEMA4D | ECM-affiliated Proteins | | X | | X | | |
| TGFB1I1 | ECM-affiliated Proteins | X | | | | | |
| ANGPTL2 | Proteoglycans | | X | | | | X |
| ASPN | Proteoglycans | X | X | X | | | X |
| BGN | Proteoglycans | X | X | X | X | X | X |
| DCN | Proteoglycans | X | X | X | X | X | X |
| HAPLN1 | Proteoglycans | | X | | | | |
| HSPG2 | Proteoglycans | X | X | X | X | X | X |
| LUM | Proteoglycans | X | X | X | X | X | X |
| OGN | Proteoglycans | X | X | | X | | X |
| PRELP | Proteoglycans | X | X | X | X | X | X |
| PRG2 | Proteoglycans | | X | | | | X |
| PRG3 | Proteoglycans | | | | | | X |
| VCAN | Proteoglycans | | X | | | X | X |
| CLU | Secreted Factors | | | | | | X |
| CRLF3 | Secreted Factors | | | | X | | |
| DCD | Secreted Factors | | | | | | X |
| DEFA1 | Secreted Factors | | | | | | X |
| EGF | Secreted Factors | X | X | | X | | |
| EGFL7 | Secreted Factors | | X | X | | | |
| FGF1 | Secreted Factors | | | | X | | |
| FGF2 | Secreted Factors | | | | | | X |
| HCFC1 | Secreted Factors | | X | | | | |
| HDGF | Secreted Factors | X | | | | | |

| | | | | | | | |
|---|---|---|---|---|---|---|---|
| HRNR | Secreted Factors | | | | | X | X |
| Il19 | Secreted Factors | X | | | | | |
| INHBE | Secreted Factors | | | | | | X |
| PF4 | Secreted Factors | | | | X | | |
| S100A1 | Secreted Factors | X | | | X | | |
| S100A6 | Secreted Factors | X | | | | | |
| S100A7 | Secreted Factors | | | | | X | |
| S100A8 | Secreted Factors | | | | X | X | X |
| S100A9 | Secreted Factors | X | | | | X | X |
| S100A10 | Secreted Factors | X | X | | X | | X |
| S100A11 | Secreted Factors | X | X | X | X | X | X |
| S100A13 | Secreted Factors | X | | | | | |
| S100A14 | Secreted Factors | | | | | | X |
| S100G | Secreted Factors | | X | | X | | |
| SFRP1 | Secreted Factors | | | | X | | |
| UMOD | Secreted Factors | X | | | | | |

**Table S3.** The comparative analysis of the matrisome composition (by categories) revealed in mouse and human kidney samples by different studies.

| | our data | McCabe 2021 | Lipp 2021 | Lui 2020 | Louzao-Martinez 2019 | Randles 2021 |
|---|---|---|---|---|---|---|
| **Extraction Method** | MCF and SE using Salt buffer, SDS and Gu-HCl | MCF | MCF | Directly solubilizing proteins in homogenized tissue by urea and sonification followed by SDS extraction | Decellularization with SDS (overnight), followed by tissue homogenisation with lysis buffer | SE using SDS and Triton |
| **Organism** | mouse | mouse | mouse | mouse | human | human |
| **Age** | 4-10 weeks | N/A | adult | 15 weeks | 55 | 15-37 |
| **Gender** | female | male | N/A | N/A | Male/female | male |
| **Strain** | C57BL/6J | C67BL/6J | C57BL/6J | C57BL/6J | N/A | N/A |
| **Size** | 3 | 3 | 3 | N/A | 13 | 3 |
| **# Total matrisome proteins** | 113 | 173 | 79 | 139 | 76 | 172 |
| **# Collagens (%)** | 16 (14%) | 22 (13%) | 18 (23%) | 16 (12%) | 12 (16%) | 21 (12%) |
| **# GPs (%)** | 28 (25%) | 55 (32%) | 38 (48%) | 41 (29%) | 33 (43%) | 53 (31%) |
| **# PGs (%)** | 7 (6%) | 10 (6%) | 6 (8%) | 6 (4%) | 6 (8%) | 11 (6%) |
| **# ECM-regulators (%)** | 36 (32%) | 57 (33%) | 9 (11%) | 46 (33%) | 11 (14%) | 47 (27%) |
| **# ECM-affiliated (%)** | 19 (17%) | 22 (13%) | 6 (8%) | 20 (14%) | 9 (12%) | 28 (16%) |
| **# Secreted factors (%)** | 7 (6%) | 7 (4%) | 2 (3%) | 10 (7%) | 5 (7%) | 12 (7%) |

Abbreviations: MCF - Millipore Compartment Fractionation; SE - Sequential Extraction Approach. N/A: not available, #:number of, GPs: ECM Glycoproteins, PGs: Proteoglycans

**Table S4.** The list of quantified matrisome proteins in Method 1 and Method 2 (all fractions) with their LFQ intensities. The list represents all matrisome proteins which had at least two LFQ intensities detected in each method. N.Q. - non quantifiable.

| Category | Gene names | Method 1 LFQ intensity | Method 2 LFQ intensity |
|---|---|---|---|
| Collagens | Col14a1 | 38691000 | 62488333.33 |
| Collagens | Col15a1 | 84182666.67 | 49110666.67 |
| Collagens | Col18a1 | 526080000 | 437766666.7 |
| Collagens | Col1a1 | 323343333.3 | 249713333.3 |
| Collagens | Col1a2 | 267973333.3 | 169754333.3 |
| Collagens | Col4a1 | 1852100000 | 863460666.7 |
| Collagens | Col4a2 | 685020000 | 250753533.3 |
| Collagens | Col4a3 | 93799666.67 | N.Q. |
| Collagens | Col4a5 | 33550000 | N.Q. |
| Collagens | Col6a1 | 550706666.7 | 253200000 |
| Collagens | Col6a2 | 594573333.3 | 330046666.7 |
| Collagens | Col6a3 | 751650000 | 309118066.7 |
| ECM Glycoproteins | Agrn | 902626666.7 | 427453333.3 |
| ECM Glycoproteins | Emilin1 | 256723333.3 | 81556666.67 |
| ECM Glycoproteins | Fbn1 | 66105333.33 | 56856666.67 |
| ECM Glycoproteins | Fga | 65870000 | 57519333.33 |
| ECM Glycoproteins | Fgg | 337723333.3 | 278483333.3 |
| ECM Glycoproteins | Fn1 | 437020000 | 215854933.3 |
| ECM Glycoproteins | Igfbp7 | 199310000 | 62936333.33 |
| ECM Glycoproteins | Lama1 | 266006666.7 | 124823333.3 |
| ECM Glycoproteins | Lama4 | 74925000 | N.Q. |
| ECM Glycoproteins | Lama5 | 382123333.3 | 556138000 |
| ECM Glycoproteins | Lamb1 | 349623333.3 | 120340566.7 |
| ECM Glycoproteins | Lamb2 | 485866666.7 | 379098000 |
| ECM Glycoproteins | Lamc1 | 680203333.3 | 258250000 |
| ECM Glycoproteins | Lgals3bp | N.Q. | 39335666.67 |
| ECM Glycoproteins | Mmrn2 | 56800000 | 47353666.67 |
| ECM Glycoproteins | Nid1 | 1628300000 | 1036566333 |
| ECM Glycoproteins | Nid2 | 784086666.7 | 292884333.3 |
| ECM Glycoproteins | Tinagl1 | 325803333.3 | 129658666.7 |
| ECM Glycoproteins | Vwa5a | N.Q. | 28909000 |
| ECM Regulators | A2m | N.Q. | 158216666.7 |
| ECM Regulators | Ace | 28259000 | 198972000 |
| ECM Regulators | Ace2 | 40415666.67 | N.Q. |
| ECM Regulators | Apoe | N.Q. | 88959000 |
| ECM Regulators | Ca15 | 291760000 | 173781666.7 |
| ECM Regulators | Cstb | N.Q. | 102790333.3 |
| ECM Regulators | Ctsa | N.Q. | 99571000 |
| ECM Regulators | Ctsb | N.Q. | 49625333.33 |
| ECM Regulators | Ctsd | N.Q. | 418300000 |
| ECM Regulators | Ctsh | N.Q. | 77440000 |
| ECM Regulators | Ctsz | N.Q. | 18227666.67 |

| | | | |
|---|---|---|---|
| ECM Regulators | Dpp4 | 378930000 | 484921000 |
| ECM Regulators | Kng1 | N.Q. | 58791666.67 |
| ECM Regulators | Mep1a | 998683333.3 | 2011577333 |
| ECM Regulators | Mep1b | 649296666.7 | 709836666.7 |
| ECM Regulators | Mme | 229546666.7 | 373354433.3 |
| ECM Regulators | Nucb1 | 11772333.33 | 97254666.67 |
| ECM Regulators | P4hb | 144053333.3 | 227476666.7 |
| ECM Regulators | Plg | 27753000 | 59242333.33 |
| ECM Regulators | Ren1 | N.Q. | 47892333.33 |
| ECM Regulators | Serpina1a | N.Q. | 204248333.3 |
| ECM Regulators | Serpina1c | N.Q. | 92267000 |
| ECM Regulators | Serpina3k | N.Q. | 474158333.3 |
| ECM Regulators | Serpinh1 | 549796666.7 | 452086666.7 |
| ECM Regulators | Sod3 | N.Q. | 241683333.3 |
| ECM Regulators | Tgm1 | N.Q. | 29695000 |
| ECM Regulators | Tgm2 | 891500000 | 339223333.3 |
| ECM-affiliated Proteins | Anxa11 | N.Q. | 60682333.33 |
| ECM-affiliated Proteins | Anxa2 | 219753333.3 | 386970000 |
| ECM-affiliated Proteins | Anxa3 | N.Q. | 51850333.33 |
| ECM-affiliated Proteins | Anxa4 | N.Q. | 34782333.33 |
| ECM-affiliated Proteins | Anxa5 | 105511333.3 | 172369333.3 |
| ECM-affiliated Proteins | Anxa6 | 54961333.33 | 133114666.7 |
| ECM-affiliated Proteins | Anxa7 | N.Q. | 20368000 |
| ECM-affiliated Proteins | C1qbp | 339133333.3 | 316836666.7 |
| ECM-affiliated Proteins | Dag1 | N.Q. | 21867666.67 |
| ECM-affiliated Proteins | Emcn | 6551366.667 | 4837190 |
| ECM-affiliated Proteins | Grn | N.Q. | 114442000 |
| ECM-affiliated Proteins | Hpx | N.Q. | 88493333.33 |
| ECM-affiliated Proteins | Lad1 | N.Q. | 12946000 |
| ECM-affiliated Proteins | Lgals1 | N.Q. | 62236666.67 |
| ECM-affiliated Proteins | Lgals3 | 24083333.33 | 131771000 |
| ECM-affiliated Proteins | Lman1 | 335713333.3 | 143900666.7 |
| ECM-affiliated Proteins | Lpl | N.Q. | 30674000 |
| ECM-affiliated Proteins | Plxnb2 | N.Q. | 88969000 |
| ECM-affiliated Proteins | Tgfb1i1 | N.Q. | 51269000 |
| Proteoglycans | Bgn | 306045666.7 | 76005766.67 |
| Proteoglycans | Dcn | 67815333.33 | N.Q. |
| Proteoglycans | Hspg2 | 2451366667 | 937683333.3 |
| Proteoglycans | Ogn | 152346666.7 | N.Q. |
| Proteoglycans | Prelp | 131366333.3 | 103712666.7 |
| Secreted Factors | S100a10 | 98618000 | 115451000 |
| Secreted Factors | S100a11 | 69156333.33 | 96207333.33 |
| Secreted Factors | S100a6 | N.Q. | 25623066.67 |
| Secreted Factors | Umod | 245273333.3 | 168239333.3 |
| Secreted Factors | Hdgf | 215473333.3 | 246296666.7 |

**Table S5.** The comparison of the LFQ intensities of the quantified matrisome proteins in the individual fraction obtained by the Method 2 and in the product of the Method 1. The list represents all matrisome proteins which had at least two LFQ intensities detected in each method. Differential analysis of LFQ intensities, in fractions of Method 2 versus Method 1, was performed only in proteins which were detected in at least two replicates of both methods. TRUE denotes significancy where p.adj (adjusted) value <0.05, FALSE is non-significant (p>0.05) and "NA" denotes to non-available. N: No; Y: Yes; F1: Fraction1; F2: Fraction 2; F3: Fraction 3.

| Method 2-F1 vs Method 1 | | | | | | | Detection in | |
|---|---|---|---|---|---|---|---|---|
| Division | Category | Gene Name | Protein ID | Fold Change | p value (adj) | Significant | Method 1 | Method 2-F1 |
| Matrisome-associated | ECM Regulators | Ace | P09470 | 2.804 | 0.015 | TRUE | Y | Y |
| Matrisome-associated | ECM Regulators | Mep1b | Q61847 | 2.678 | 0.008 | TRUE | Y | Y |
| Matrisome-associated | ECM Regulators | Ctsa | G3X8T3 | 2.406 | 0.045 | TRUE | Y | Y |
| Core matrisome | Proteoglycans | Hspg2 | E9PZ16 | -2.240 | 0.021 | TRUE | Y | Y |
| Core matrisome | ECM Glycoproteins | Fn1 | P11276 | -2.284 | 0.024 | TRUE | Y | Y |
| Core matrisome | Collagens | Col18a1 | P39061 | -2.405 | 0.026 | TRUE | Y | Y |
| Core matrisome | ECM Glycoproteins | Emilin1 | Q99K41 | -2.501 | 0.015 | TRUE | Y | Y |
| Matrisome-associated | ECM Regulators | Tgm2 | P21981 | -2.942 | 0.004 | TRUE | Y | Y |
| Core matrisome | Proteoglycans | Prelp | Q9JK53 | -3.141 | 0.018 | TRUE | Y | Y |
| Matrisome-associated | ECM Regulators | Serpinh1 | P19324 | -3.408 | 0.027 | TRUE | Y | Y |
| Core matrisome | ECM Glycoproteins | Agrn | M0QWP1 | -3.592 | 0.002 | TRUE | Y | Y |
| Core matrisome | ECM Glycoproteins | Tinagl1 | Q99JR5 | -4.266 | 0.007 | TRUE | Y | Y |
| Core matrisome | ECM Glycoproteins | Nid2 | O88322 | -4.752 | 0.003 | TRUE | Y | Y |
| Core matrisome | ECM Glycoproteins | Nid1 | P10493 | -5.282 | 0.003 | TRUE | Y | Y |
| Core matrisome | Collagens | Col4a3 | Q9QZS0 | 3.927 | 0.141 | FALSE | Y | Y |
| Matrisome-associated | ECM Regulators | Ctsb | P10605 | 3.578 | 0.185 | FALSE | Y | Y |
| Core matrisome | Collagens | Col4a5 | Q63ZW6 | 2.402 | 0.247 | FALSE | Y | Y |
| Matrisome-associated | ECM Regulators | Mme | Q61391 | 1.785 | 0.052 | FALSE | Y | Y |
| Matrisome-associated | ECM Regulators | Ace2 | Q8R0I0 | 1.722 | 0.096 | FALSE | Y | Y |
| Core matrisome | Collagens | Col1a1 | P11087 | 1.563 | 0.248 | FALSE | Y | Y |
| Core matrisome | ECM Glycoproteins | Fbn1 | Q61554 | 1.538 | 0.155 | FALSE | Y | Y |
| Core matrisome | ECM Glycoproteins | Lama5 | Q61001 | 1.399 | 0.284 | FALSE | Y | Y |
| Core matrisome | Collagens | Col4a1 | P02463 | 1.374 | 0.276 | FALSE | Y | Y |
| Matrisome-associated | ECM Regulators | Tgm1 | Q9JLF6 | 1.336 | 0.367 | FALSE | Y | Y |
| Matrisome-associated | ECM-affiliated Proteins | Lman1 | Q9D0F3 | 1.335 | 0.333 | FALSE | Y | Y |
| Matrisome-associated | ECM Regulators | Dpp4 | P28843 | 1.321 | 0.284 | FALSE | Y | Y |
| Core matrisome | ECM Glycoproteins | Fgg | Q8VCM7 | 1.287 | 0.418 | FALSE | Y | Y |
| Matrisome-associated | ECM Regulators | Mep1a | P28825 | 1.278 | 0.468 | FALSE | Y | Y |
| Matrisome-associated | ECM Regulators | Ca15 | Q99N23 | 1.259 | 0.630 | FALSE | Y | Y |
| Core matrisome | Collagens | Col1a2 | Q01149 | 1.100 | 0.798 | FALSE | Y | Y |
| Core matrisome | Collagens | Col6a2 | Q02788 | -1.025 | 0.942 | FALSE | Y | Y |
| Matrisome-associated | ECM-affiliated Proteins | Anxa6 | P14824 | -1.030 | 0.947 | FALSE | Y | Y |
| Core matrisome | ECM Glycoproteins | Lamc1 | F8VQJ3 | -1.082 | 0.704 | FALSE | Y | Y |
| Core matrisome | Collagens | Col4a2 | P08122 | -1.105 | 0.742 | FALSE | Y | Y |

| | | | | | | | | |
|---|---|---|---|---|---|---|---|---|
| Core matrisome | ECM Glycoproteins | Lama1 | P19137 | -1.129 | 0.592 | FALSE | Y | Y |
| Matrisome-associated | ECM-affiliated Proteins | Lpl | P11152 | -1.166 | 0.803 | FALSE | Y | Y |
| Core matrisome | ECM Glycoproteins | Lamb2 | Q61292 | -1.175 | 0.461 | FALSE | Y | Y |
| Core matrisome | Collagens | Col6a1 | Q04857 | -1.178 | 0.548 | FALSE | Y | Y |
| Core matrisome | Collagens | Col14a1 | Q80X19 | -1.182 | 0.827 | FALSE | Y | Y |
| Matrisome-associated | ECM-affiliated Proteins | Anxa5 | P48036 | -1.186 | 0.593 | FALSE | Y | Y |
| Matrisome-associated | Secreted Factors | S100a11 | P50543 | -1.221 | 0.569 | FALSE | Y | Y |
| Matrisome-associated | Secreted Factors | Umod | Q91X17 | -1.230 | 0.361 | FALSE | Y | Y |
| Matrisome-associated | ECM-affiliated Proteins | Anxa2 | P07356 | -1.251 | 0.687 | FALSE | Y | Y |
| Core matrisome | Collagens | Col6a3 | E9PWQ3 | -1.275 | 0.301 | FALSE | Y | Y |
| Core matrisome | ECM Glycoproteins | Lamb1 | P02469 | -1.381 | 0.220 | FALSE | Y | Y |
| Core matrisome | ECM Glycoproteins | Fga | E9PV24 | -1.416 | 0.373 | FALSE | Y | Y |
| Matrisome-associated | ECM Regulators | P4hb | P09103 | -1.639 | 0.330 | FALSE | Y | Y |
| Core matrisome | Collagens | Col15a1 | A2AJY2 | -1.930 | 0.170 | FALSE | Y | Y |
| Matrisome-associated | Secreted Factors | S100a10 | P08207 | -3.236 | 0.051 | FALSE | Y | Y |
| Core matrisome | Collagens | Col4a4 | Q9QZR9 | NA | NA | NA | N | Y |
| Core matrisome | ECM Glycoproteins | Ltbp4 | Q8K4G1 | NA | NA | NA | N | Y |
| Core matrisome | ECM Glycoproteins | Grn | P28798 | NA | NA | NA | N | y |
| Core matrisome | ECM Glycoproteins | Npnt | Q91V88 | NA | NA | NA | N | Y |
| Matrisome-associated | ECM Regulators | Plg | P20918 | NA | NA | NA | N | Y |
| Matrisome-associated | ECM Regulators | Nucb1 | Q02819 | NA | NA | NA | N | Y |
| Core matrisome | ECM Glycoproteins | Igfbp7 | E9Q5D9 | NA | NA | NA | Y | N |
| Core matrisome | ECM Glycoproteins | Lama4 | P97927 | NA | NA | NA | Y | N |
| Core matrisome | Proteoglycans | Bgn | P28653 | NA | NA | NA | Y | N |
| Core matrisome | Proteoglycans | Ogn | Q62000 | NA | NA | NA | Y | N |
| Core matrisome | Proteoglycans | Dcn | P28654 | NA | NA | NA | Y | N |
| Matrisome-associated | ECM-affiliated Proteins | C1qbp | O35658 | NA | NA | NA | Y | N |
| Matrisome-associated | ECM-affiliated Proteins | Lad1 | P57016 | NA | NA | NA | Y | N |
| Matrisome-associated | ECM-affiliated Proteins | Tgfb1i1 | Q62219 | NA | NA | NA | Y | N |
| **Method 2-F2 vs Method 1** | | | | | | | **Detection in** | |
| **Division** | **Category** | **Gene Name** | **Protein IDs** | **Fold Change** | **p value (adj)** | **Significant** | **Method 1** | **Method 2-F2** |
| Matrisome-associated | ECM Regulators | Serpina1a | P07758 | 49.867 | 0.000 | TRUE | Y | Y |
| Matrisome-associated | ECM Regulators | A2m | Q61838 | 20.112 | 0.000 | TRUE | Y | Y |
| Matrisome-associated | ECM-affiliated Proteins | Lgals1 | P16045 | 4.659 | 0.001 | TRUE | Y | Y |
| Matrisome-associated | ECM-affiliated Proteins | Anxa6 | P14824 | 3.605 | 0.009 | TRUE | Y | Y |
| Matrisome-associated | ECM Regulators | Ctsa | G3X8T3 | 3.227 | 0.009 | TRUE | Y | Y |
| Matrisome-associated | ECM-affiliated Proteins | Anxa5 | P48036 | 2.694 | 0.018 | TRUE | Y | Y |
| Matrisome-associated | ECM Regulators | Ctsd | P18242 | 2.567 | 0.015 | TRUE | Y | Y |
| Matrisome-associated | ECM-affiliated Proteins | Anxa2 | P07356 | 2.514 | 0.025 | TRUE | Y | Y |

| | | | | | | | | |
|---|---|---|---|---|---|---|---|---|
| Matrisome-associated | Secreted Factors | S100a11 | P50543 | 2.297 | 0.009 | TRUE | Y | Y |
| Matrisome-associated | Secreted Factors | Hdgf | P51859 | 2.056 | 0.029 | TRUE | Y | Y |
| Core matrisome | ECM Glycoproteins | Fgg | Q8VCM7 | -2.585 | 0.016 | TRUE | Y | Y |
| Core matrisome | ECM Glycoproteins | Igfbp7 | E9Q5D9 | -2.751 | 0.003 | TRUE | Y | Y |
| Core matrisome | ECM Glycoproteins | Fga | E9PV24 | -2.908 | 0.002 | TRUE | Y | Y |
| Matrisome-associated | ECM Regulators | Serpinh1 | P19324 | -4.347 | 0.001 | TRUE | Y | Y |
| Matrisome-associated | ECM Regulators | Tgm2 | P21981 | -6.190 | 0.000 | TRUE | Y | Y |
| Matrisome-associated | Secreted Factors | Umod | Q91X17 | -8.877 | 0.000 | TRUE | Y | Y |
| Matrisome-associated | ECM Regulators | Mep1a | P28825 | -13.361 | 0.000 | TRUE | Y | Y |
| Core matrisome | Proteoglycans | Hspg2 | E9PZ16 | -28.443 | 0.000 | TRUE | Y | Y |
| Core matrisome | ECM Glycoproteins | Lamb1 | P02469 | -33.359 | 0.000 | TRUE | Y | Y |
| Core matrisome | ECM Glycoproteins | Nid1 | P10493 | -64.445 | 0.000 | TRUE | Y | Y |
| Matrisome-associated | ECM Regulators | Sod3 | O09164 | 5.579 | 0.077 | FALSE | Y | Y |
| Core matrisome | Collagens | Col14a1 | Q80X19 | 2.114 | 0.264 | FALSE | Y | Y |
| Core matrisome | ECM Glycoproteins | Mmrn2 | A6H6E2 | 1.691 | 0.491 | FALSE | Y | Y |
| Matrisome-associated | ECM-affiliated Proteins | Lad1 | P57016 | 1.268 | 0.334 | FALSE | Y | Y |
| Matrisome-associated | Secreted Factors | S100a10 | P08207 | 1.072 | 0.814 | FALSE | Y | Y |
| Matrisome-associated | ECM Regulators | P4hb | P09103 | 1.017 | 0.951 | FALSE | Y | Y |
| Matrisome-associated | ECM-affiliated Proteins | C1qbp | O35658 | -1.852 | 0.086 | FALSE | Y | Y |
| Matrisome-associated | ECM Regulators | Ctsz | Q9WUU7 | NA | NA | NA | N | Y |
| Matrisome-associated | ECM Regulators | Apoe | P08226 | NA | NA | NA | N | Y |
| Matrisome-associated | ECM Regulators | Ren1 | P06281 | NA | NA | NA | N | Y |
| Matrisome-associated | ECM Regulators | Hrg | Q9ESB3 | NA | NA | NA | N | Y |
| Matrisome-associated | ECM Regulators | Serpina1b | P22599 | NA | NA | NA | N | Y |
| Matrisome-associated | ECM Regulators | Ctsh | P49935 | NA | NA | NA | N | Y |
| Matrisome-associated | ECM Regulators | Kng1 | O08677 | NA | NA | NA | N | Y |
| Matrisome-associated | ECM Regulators | Cstb | Q62426 | NA | NA | NA | N | Y |
| Matrisome-associated | ECM Regulators | Ctsb | P10605 | NA | NA | NA | N | Y |
| Matrisome-associated | ECM Regulators | Serpina3k | A0A0R4J0I1 | NA | NA | NA | N | Y |
| Matrisome-associated | ECM-affiliated Proteins | Anxa11 | P97384 | NA | NA | NA | N | Y |
| Matrisome-associated | ECM-affiliated Proteins | Anxa7 | Q07076 | NA | NA | NA | N | Y |
| Matrisome-associated | ECM-affiliated Proteins | Anxa3 | O35639 | NA | NA | NA | N | Y |
| Matrisome-associated | ECM-affiliated Proteins | Anxa4 | P97429 | NA | NA | NA | N | Y |
| Matrisome-associated | ECM-affiliated Proteins | Hpx | Q91X72 | NA | NA | NA | N | Y |
| Matrisome-associated | Secreted Factors | S100a6 | P14069 | NA | NA | NA | N | Y |
| Core matrisome | ECM Glycoproteins | Grn | P28798 | NA | NA | NA | N | Y |
| Matrisome-associated | ECM Regulators | Plg | P20918 | NA | NA | NA | N | Y |

| | | | | | | | | |
|---|---|---|---|---|---|---|---|---|
| Matrisome-associated | ECM Regulators | Arsb | A0A0R4J138 | NA | NA | NA | N | Y |
| Matrisome-associated | ECM Regulators | F2 | P19221 | NA | NA | NA | N | Y |
| Matrisome-associated | ECM Regulators | Galns | Q8CC47 | NA | NA | NA | N | Y |
| Core matrisome | Collagens | Col6a3 | E9PWQ3 | NA | NA | NA | Y | N |
| Core matrisome | ECM Glycoproteins | Fn1 | P11276 | NA | NA | NA | Y | N |
| Core matrisome | Collagens | Col4a1 | P02463 | NA | NA | NA | Y | N |
| Core matrisome | Collagens | Col4a2 | P08122 | NA | NA | NA | Y | N |
| Core matrisome | Collagens | Col6a2 | Q02788 | NA | NA | NA | Y | N |
| Core matrisome | Collagens | Col1a1 | P11087 | NA | NA | NA | Y | N |
| Core matrisome | Collagens | Col1a2 | Q01149 | NA | NA | NA | Y | N |
| Core matrisome | Collagens | Col6a1 | Q04857 | NA | NA | NA | Y | N |
| Core matrisome | Collagens | Col18a1 | P39061 | NA | NA | NA | Y | N |
| Core matrisome | Collagens | Col15a1 | A2AJY2 | NA | NA | NA | Y | N |
| Core matrisome | ECM Glycoproteins | Nid2 | O88322 | NA | NA | NA | Y | N |
| Core matrisome | ECM Glycoproteins | Agrn | M0QWP1 | NA | NA | NA | Y | N |
| Core matrisome | ECM Glycoproteins | Lamc1 | F8VQJ3 | NA | NA | NA | Y | N |
| Core matrisome | ECM Glycoproteins | Lama5 | Q61001 | NA | NA | NA | Y | N |
| Core matrisome | ECM Glycoproteins | Tinagl1 | Q99JR5 | NA | NA | NA | Y | N |
| Core matrisome | ECM Glycoproteins | Lama1 | P19137 | NA | NA | NA | Y | N |
| Core matrisome | ECM Glycoproteins | Lamb2 | Q61292 | NA | NA | NA | Y | N |
| Core matrisome | ECM Glycoproteins | Emilin1 | Q99K41 | NA | NA | NA | Y | N |
| Core matrisome | ECM Glycoproteins | Fbn1 | Q61554 | NA | NA | NA | Y | N |
| Core matrisome | ECM Glycoproteins | Lama4 | P97927 | NA | NA | NA | Y | N |
| Core matrisome | Proteoglycans | Bgn | P28653 | NA | NA | NA | Y | N |
| Core matrisome | Proteoglycans | Prelp | Q9JK53 | NA | NA | NA | Y | N |
| Core matrisome | Proteoglycans | Ogn | Q62000 | NA | NA | NA | Y | N |
| Core matrisome | Proteoglycans | Dcn | P28654 | NA | NA | NA | Y | N |
| Matrisome-associated | ECM Regulators | Mep1b | Q61847 | NA | NA | NA | Y | N |
| Matrisome-associated | ECM Regulators | Dpp4 | P28843 | NA | NA | NA | Y | N |
| Matrisome-associated | ECM Regulators | Mme | Q61391 | NA | NA | NA | Y | N |
| Matrisome-associated | ECM Regulators | Ca15 | Q99N23 | NA | NA | NA | Y | N |
| Matrisome-associated | ECM Regulators | Ace | P09470 | NA | NA | NA | Y | N |
| Matrisome-associated | ECM Regulators | Ace2 | Q8R0I0 | NA | NA | NA | Y | N |
| Matrisome-associated | ECM-affiliated Proteins | Lman1 | Q9D0F3 | NA | NA | NA | Y | N |
| Core matrisome | Collagens | Col4a3 | Q9QZS0 | NA | NA | NA | Y | N |
| Core matrisome | Collagens | Col4a5 | Q63ZW6 | NA | NA | NA | Y | N |
| Core matrisome | ECM Glycoproteins | Vwa5a | Q99KC8 | NA | NA | NA | Y | N |
| **Method 2-F3 vs Method 1** | | | | | | | **Detection in** | |
| **Division** | **Category** | **Gene Name** | **Protein IDs** | **Fold Change** | **p value (adj)** | **Significant** | **Method 1** | **Method 2-F3** |
| Matrisome-associated | ECM-affiliated Proteins | Lgals1 | P16045 | 6.498 | 0.005 | TRUE | Y | Y |
| Matrisome-associated | ECM Regulators | Mep1a | P28825 | 3.758 | 0.005 | TRUE | Y | Y |
| Matrisome-associated | ECM Regulators | Plg | P20918 | 3.160 | 0.034 | TRUE | Y | Y |

| | | | | | | | | |
|---|---|---|---|---|---|---|---|---|
| Matrisome-associated | ECM Regulators | P4hb | P09103 | 2.099 | 0.017 | TRUE | Y | Y |
| Matrisome-associated | ECM Regulators | Ace | P09470 | -2.189 | 0.039 | TRUE | Y | Y |
| Matrisome-associated | ECM Regulators | Tgm2 | P21981 | -2.189 | 0.013 | TRUE | Y | Y |
| Core matrisome | Proteoglycans | Ogn | Q62000 | -2.204 | 0.013 | TRUE | Y | Y |
| Matrisome-associated | ECM Regulators | Mep1b | Q61847 | -2.297 | 0.013 | TRUE | Y | Y |
| Core matrisome | ECM Glycoproteins | Igfbp7 | E9Q5D9 | -2.532 | 0.012 | TRUE | Y | Y |
| Core matrisome | ECM Glycoproteins | Lama1 | P19137 | -2.639 | 0.012 | TRUE | Y | Y |
| Matrisome-associated | Secreted Factors | S100a11 | P50543 | -3.074 | 0.036 | TRUE | Y | Y |
| Core matrisome | ECM Glycoproteins | Lama5 | Q61001 | -3.095 | 0.017 | TRUE | Y | Y |
| Core matrisome | ECM Glycoproteins | Fga | E9PV24 | -3.138 | 0.026 | TRUE | Y | Y |
| Core matrisome | Collagens | Col1a2 | Q01149 | -3.160 | 0.008 | TRUE | Y | Y |
| Core matrisome | Collagens | Col6a1 | Q04857 | -3.204 | 0.007 | TRUE | Y | Y |
| Core matrisome | ECM Glycoproteins | Emilin1 | Q99K41 | -3.204 | 0.004 | TRUE | Y | Y |
| Core matrisome | Collagens | Col6a3 | E9PWQ3 | -3.531 | 0.005 | TRUE | Y | Y |
| Core matrisome | Collagens | Col6a2 | Q02788 | -3.681 | 0.036 | TRUE | Y | Y |
| Matrisome-associated | ECM Regulators | Mme | Q61391 | -4.112 | 0.004 | TRUE | Y | Y |
| Core matrisome | ECM Glycoproteins | Lamb2 | Q61292 | -4.469 | 0.001 | TRUE | Y | Y |
| Matrisome-associated | ECM Regulators | Dpp4 | P28843 | -5.352 | 0.001 | TRUE | Y | Y |
| Core matrisome | ECM Glycoproteins | Lamc1 | F8VQJ3 | -5.389 | 0.002 | TRUE | Y | Y |
| Core matrisome | ECM Glycoproteins | Lamb1 | P02469 | -6.105 | 0.004 | TRUE | Y | Y |
| Matrisome-associated | ECM Regulators | Ca15 | Q99N23 | -9.714 | 0.013 | TRUE | Y | Y |
| Core matrisome | Collagens | Col4a1 | P02463 | -52.710 | 0.001 | TRUE | Y | Y |
| Matrisome-associated | ECM-affiliated Proteins | Lgals3 | P16110 | 3.074 | 0.054 | FALSE | Y | Y |
| Matrisome-associated | ECM Regulators | Serpina1c | A0A0R4J0X5 | 1.968 | 0.024 | FALSE | Y | Y |
| Matrisome-associated | ECM-affiliated Proteins | C1qbp | O35658 | 1.889 | 0.041 | FALSE | Y | Y |
| Matrisome-associated | ECM Regulators | Nucb1 | Q02819 | 1.620 | 0.086 | FALSE | Y | Y |
| Matrisome-associated | ECM Regulators | Serpinh1 | P19324 | 1.474 | 0.104 | FALSE | Y | Y |
| Matrisome-associated | ECM-affiliated Proteins | Plxnb2 | B2RXS4 | 1.402 | 0.370 | FALSE | Y | Y |
| Core matrisome | Collagens | Col18a1 | P39061 | 1.396 | 0.112 | FALSE | Y | Y |
| Core matrisome | Proteoglycans | Lum | P51885 | 1.352 | 0.279 | FALSE | Y | Y |
| Core matrisome | Proteoglycans | Prelp | Q9JK53 | 1.310 | 0.206 | FALSE | Y | Y |
| Matrisome-associated | ECM-affiliated Proteins | Anxa2 | P07356 | 1.198 | 0.689 | FALSE | Y | Y |
| Matrisome-associated | Secreted Factors | Hdgf | P51859 | 1.078 | 0.824 | FALSE | Y | Y |
| Matrisome-associated | ECM-affiliated Proteins | Anxa6 | P14824 | 1.067 | 0.829 | FALSE | Y | Y |
| Core matrisome | Collagens | Col15a1 | A2AJY2 | 1.053 | 0.849 | FALSE | Y | Y |
| Matrisome-associated | Secreted Factors | S100a10 | P08207 | 1.047 | 0.884 | FALSE | Y | Y |
| Core matrisome | ECM Glycoproteins | Agrn | M0QWP1 | 1.033 | 0.879 | FALSE | Y | Y |
| Matrisome-associated | ECM-affiliated Proteins | Lad1 | P57016 | 1.013 | 0.954 | FALSE | Y | Y |
| Core matrisome | Collagens | Col14a1 | Q80X19 | -1.059 | 0.923 | FALSE | Y | Y |
| Matrisome-associated | Secreted Factors | Umod | Q91X17 | -1.094 | 0.696 | FALSE | Y | Y |

| | | | | | | | | |
|---|---|---|---|---|---|---|---|---|
| Core matrisome | Proteoglycans | Bgn | P28653 | -1.194 | 0.565 | FALSE | Y | Y |
| Core matrisome | ECM Glycoproteins | Nid1 | P10493 | -1.228 | 0.339 | FALSE | Y | Y |
| Core matrisome | Proteoglycans | Dcn | P28654 | -1.244 | 0.585 | FALSE | Y | Y |
| Core matrisome | ECM Glycoproteins | Fn1 | P11276 | -1.303 | 0.244 | FALSE | Y | Y |
| Core matrisome | ECM Glycoproteins | Tinagl1 | Q99JR5 | -1.304 | 0.268 | FALSE | Y | Y |
| Core matrisome | ECM Glycoproteins | Nid2 | O88322 | -1.482 | 0.098 | FALSE | Y | Y |
| Core matrisome | Proteoglycans | Hspg2 | E9PZ16 | -1.515 | 0.086 | FALSE | Y | Y |
| Matrisome-associated | ECM-affiliated Proteins | Anxa5 | P48036 | -1.696 | 0.038 | FALSE | Y | Y |
| Core matrisome | Collagens | Col1a1 | P11087 | -2.014 | 0.052 | FALSE | Y | Y |
| Core matrisome | ECM Glycoproteins | Dpt | Q9QZZ6 | NA | NA | NA | N | Y |
| Core matrisome | ECM Glycoproteins | Grn | P28798 | NA | NA | NA | N | Y |
| Matrisome-associated | ECM Regulators | Ctsa | G3X8T3 | NA | NA | NA | N | Y |
| Matrisome-associated | ECM Regulators | Apoe | P08226 | NA | NA | NA | N | Y |
| Matrisome-associated | ECM-affiliated Proteins | Hpx | Q91X72 | NA | NA | NA | N | Y |
| Core matrisome | ECM Glycoproteins | Lgals3bp | Q07797 | NA | NA | NA | N | Y |
| Matrisome-associated | ECM Regulators | A2m | Q61838 | NA | NA | NA | N | Y |
| Matrisome-associated | ECM-affiliated Proteins | Tgfb1i1 | Q62219 | NA | NA | NA | N | Y |
| Core matrisome | Collagens | Col4a2 | P08122 | NA | NA | NA | Y | N |
| Core matrisome | ECM Glycoproteins | Fbn1 | Q61554 | NA | NA | NA | Y | N |
| Core matrisome | ECM Glycoproteins | Lama4 | P97927 | NA | NA | NA | Y | N |
| Matrisome-associated | ECM Regulators | Ace2 | Q8R0I0 | NA | NA | NA | Y | N |
| Matrisome-associated | ECM-affiliated Proteins | Lman1 | Q9D0F3 | NA | NA | NA | Y | N |
| Core matrisome | Collagens | Col4a3 | Q9QZS0 | NA | NA | NA | Y | N |
| Core matrisome | Collagens | Col4a5 | Q63ZW6 | NA | NA | NA | Y | N |
| Core matrisome | ECM Glycoproteins | Mmrn2 | A6H6E2 | NA | NA | NA | Y | N |
| Matrisome-associated | ECM-affiliated Proteins | Lpl | P11152 | NA | NA | NA | Y | N |

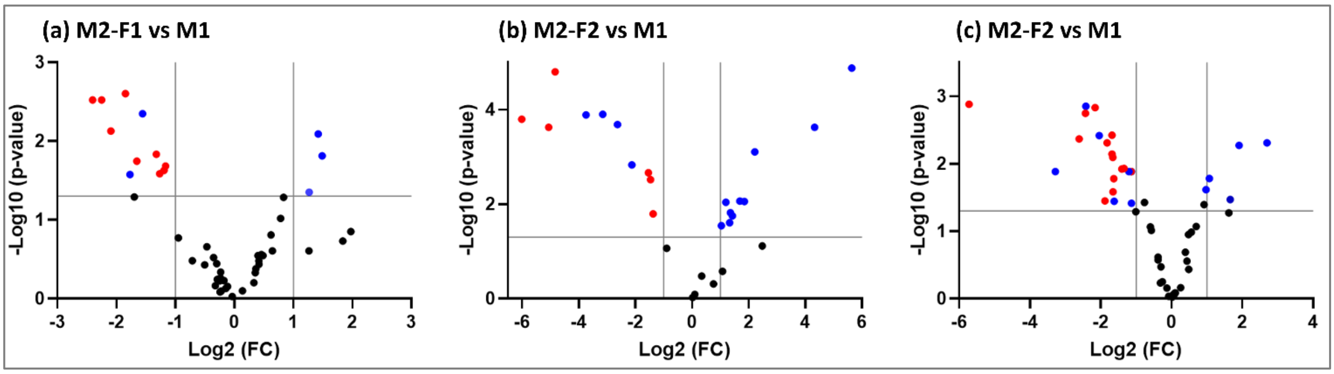


**Figure S2 .** Quantitative comparison of amount of protein extraction levels between Method 2 fractions and Method 1. Volcano plots represent significant differences in protein abundances (FC > 2, p-value < 0.05) between **(a)** Method 2 (faction 1) vs Method 1, **(b)** Method 2 (faction 2) vs Method 1 and **(c)** Method 2 (fraction 3) vs Method 1. Top-right hand sides of volcano plots are proteins that were significantly higher in Method 2 fractions. Top-left hand sides are proteins that were significantly higher in Method 1. Blue dots indicate Matrisome-associated proteins and red dots indicate Core matrisome proteins that were significantly higher in either Method 2 fractions or Method 1.
